# Multicolor structured illumination microscopy and quantitative control of polychromatic coherent light with a digital micromirror device

**DOI:** 10.1101/2020.07.27.223941

**Authors:** Peter T. Brown, Rory Kruithoff, Gregory J. Seedorf, Douglas P. Shepherd

**Affiliations:** Department of Physics and Center for Biological Physics, Arizona State University, Tempe, AZ 85287, USA; Department of Pediatrics and Pediatric Heart Lung Center, University of Colorado Anschutz Medical Campus, Aurora, CO 80045, USA

## Abstract

Structured illumination microscopy (SIM) is a broadly applicable super-resolution microscopy technique which does not impose photophysics requirements on fluorescent samples. Multicolor SIM implementations typically rely on liquid crystal on silicon (LCoS) spatial light modulators (SLM’s) for precise patterning of the excitation light, but digital micromirror devices (DMD’s) are a promising alternative, owing to their lower cost, increased imaging rate, and simplified experimental timings. Given these advantages, why do existing DMD SIM implementations either rely on incoherent projection, resulting in an order of magnitude lower signal-to-noise, or utilize coherent light at only a single wavelength? The primary obstacle to realizing a multicolor coherent DMD SIM microscope is the lack of an efficient approach for dealing with the blazed grating effect. To address this challenge, we developed quantitative tools applicable to a single DMD acting as a polychromatic diffractive optic. These include a closed form solution of the blaze and diffraction conditions, a forward model of DMD diffraction, and a forward model of coherent pattern projection. We applied these to identify experimentally feasible configurations using a single DMD as a polychromatic diffractive optic for combinations of three and four common fluorophore wavelengths. Based on these advances, we constructed a DMD SIM microscope for coherent light which we used to validate these models, develop a high-resolution optical transfer function measurement technique, and demonstrate SIM resolution enhancement for calibration samples, fixed cells, and live cells. This low-cost setup opens the door to applying DMD’s in polychromatic applications which were previously restricted to LCoS SLM’s.

## 1: Introduction

The development of wavefront control using spatial light modulators (SLM’s) has enabled a variety of techniques relevant to the study of biological and atomic systems, including quantitative phase imaging (1,2), optical trapping (3–5), adaptive optics (6), production of arbitrary dynamic optical potentials, and quantum gas microscopy (7, 8). SLM’s are highly reconfigurable, which allows experiments to be modular and take advantage of dynamic light patterns. They are also highly reproducible and offer microscopic control of wavefronts, which allows precise calibration of optical systems and enhances quantitative modeling of experiments. Due to this broad applicability, advancements in optical methodology for SLM’s have had an immediate impact for many fields of quantitative imaging.

The study of biological regulation at the molecular level is one field that has greatly benefited from advancements in optical methodologies. For example, the development of super-resolution (SR) microscopy has allowed optical study of biological systems below the diffraction limit, on the 1 nm-200 nm scale. Despite the promise of sub-diffraction studies of molecular interactions, SR techniques have yielded limited insight into the temporal dynamics of molecular regulation within living cells and larger systems. One barrier is that all SR methods impose a trade-off between imaging speed, resolution, sample preparation, and fluorophore photophysics. For example, single molecule localization microscopy (SMLM) trades temporal resolution for spatial resolution, requiring fixed samples and repeated imaging of fluorophores with specific photophysics (9, 10). Computational methods that infer SR information from fluctuating signals relax the above requirements, but require acquisition, processing, and merging of many imaging frames (11, 12). Stimulated emission depletion (STED) requires a high intensity depletion beam, careful alignment of the depletion and excitation beams, specific fluorophores, and raster scans to build an image (13, 14). MINFLUX and MINSTED lower the total intensity incident on the sample as compared to STED, but still requires raster scanning (15, 16). Methods such as MoNaLISA parallelize the application of depletion and saturation to imaging, but require precise 3D pattern projection and specific fluorophore photophysics (17, 18). In contrast to the above methods, structured illumination microscopy (SIM) does not impose specific sample requirements. The trade-off for this flexibility is a modest increase in high frequency content for linear SIM, requirement for high-quality control of the excitation light, and potentially slow imaging rates. New quantitative approaches that both simplify the construction and increase the speed of multicolor SIM would provide researchers with a flexible platform to measure temporal dynamics below the classical diffraction limit using standard sample preparations.

Here, we briefly outline the working principles, current state-of-the-art, and limitations of existing SIM instruments. SIM obtains SR information by projecting a sinusoidal illumination pattern on the sample and relying on the Moiré effect to down-mix high-frequency sample information below the diffraction limit. It was first proposed as an SR technique in (19, 20), but a similar technique was previously exploited for optical sectioning (OS) (21, 22). Early implementations achieved SR in the lateral direction only (2D-SIM) (23,24), while later approaches also enhanced the axial resolution (3D-SIM) (25–28). Initial experiments utilized diffraction gratings to produce SIM patterns and required both separate paths for multiple colors and physical translation and rotation, which severely limited their speed (27). Various advances in LCoS SLM’s, sCMOS cameras, and GPU’s have enabled much faster 2D (29–34) and 3D SIM (35), and multicolor SIM with a single optical path (36). Recent experiments have exploited SIM techniques to increase the precision of localization microscopy (37, 38). The quality of the obtained SIM reconstructions is highly dependent on the modulation depth of the projected patterns and any other deviations from the ideal optical transfer function used to weight the different frequency components. As the complexity of the desired result increases from obtaining optical sectioning to multicolor 3D-SIM, so does the required fidelity in the final projected patterns. This has led to wide spread adoption of LCoS SLM’s as a diffractive optic for SIM, despite their drawbacks which include cost, complex experimental timing, and relatively slow speed.

Digital micromirror devices (DMD’s) are a promising alternative to LCoS SLM’s for a variety of wavefront shaping tasks. DMD’s offer several advantages, including a factor of 2–10 lower cost, less experimental timing complexity, and a factor of 5–10 imaging rate increase (39–41). DMD’s can reach frame rates of up to 30 kHz, exceeding the rates achievable in ferroelectric and nematic LCoS SLM’s by factors of ~5 and ~10 respectively. Unlike ferroelectric SLM’s, DMD’s do not require the pattern to be inverted every ~100 ms. Finally, the amplitude-only modulation characteristic of the DMD allows fast, well-defined diversion of the illumination beam from the sample, making an additional fast shutter unnecessary. However, DMD use is currently limited by a computationally expensive forward model (40) to evaluate the blazed grating effect, which enhances the diffraction efficiency into a single order but also imposes severe restrictions on multicolor operation. To date, DMD-SIM approaches have used incoherent projection (42) or one coherent wavelength (40, 41, 43). Incoherent projection SIM at best provides an order of magnitude lower signal-to-noise ratio, leading to inferior experimental resolution, despite previously reported erroneous resolution measurements (42, 44) (Supplemental Note 10).

In this work, we overcome these difficulties to realize a flexible, three-color DMD microscope using coherent light. We achieve this by creating analytic forward models of both diffraction from the DMD and SIM pattern projection which limit computation time by identifying the discrete set of diffracted frequencies generated by a given pattern. Next, we harness these models to design an experimental optical configuration that satisfies the unique requirements the DMD imposes when used as a polychromatic diffractive optic. Finally, we employ this apparatus in three applications: (1) we validate these forward models by performing a detailed comparison between the positions and intensities of diffraction peaks generated by hundreds of SIM patterns, (2) we develop a calibration routine to directly map the optical transfer function (OTF) of the system without the need to estimate it from sub-diffraction limited fluorescent microspheres, and (3) we realize three wavelength, 2D linear SIM of calibration samples, fixed cells, and live cells. These applications illustrate the potential of this framework to enhance rational and quantitative design of DMD based instruments.

## 2: Results

Leveraging the advantages of the DMD as a polychromatic diffractive optic requires a tractable forward model of diffraction from the displayed DMD patterns. To this end, we developed an analytic solution of the combined blaze and diffraction condition for arbitrary wavelengths and incidence angles (Supplemental Note 1) and derived analytical forward models of DMD diffraction and pattern projection (see Materials and Methods) which we validated experimentally (Supplemental Notes 2-3). This provides an alternative framework to a previously published numerical simulation approach (40). We then applied this theoretical approach to design a DMD microscope for three-color coherent light operation. With this instrument, we developed a technique to measure the system optical transfer function and realized three-color SIM imaging at ~470 nm, 532 nm, and 635 nm over a 100 μm × 90 μm field of view using a 100 × fluorite objective with na = 1.3. This field of view is at least four times as large as typical DMD SIM (40, 41) and LCoS SLM instruments (34, 45). We validated this instrument’s SIM capabilities by imaging a variety of samples, including an Argo-SIM calibration sample, which shows our instrument produces SR information in all spatial directions and achieves resolution near the theoretical limit, and fixed and live cells, which demonstrate two- and three-color imaging with SR enhancement in biological systems.

To support our instrument and disseminate our quantitative modeling tools, we created a Python-based software suite for computing forward models of DMD diffraction and pattern projection, DMD pattern design, OTF measurement analysis, simulation of SIM imaging given a ground truth structure, and 2D SIM reconstruction. Our SIM reconstruction algorithm is based on published work (46), with enhancements based on the ability to precisely calibrate fringe projection and direct measurement of the OTF (Supplemental Note 7). We compared reconstruction results with FairSIM (Supplemental Note 8) and explored their behavior versus SNR under a variety of conditions through simulation (Supplemental Note 9). The software suite is available on GitHub, https://github.com/QI2lab/mcSlM.

### 2.1. Optical transfer function determination

As a first demonstration of the advantages of our DMD and SIM pattern forward models for quantitative imaging, we apply them to directly measure the optical transfer function by projecting a sequence of SIM patterns, including all diffraction orders, on a fluorescent sample and comparing the strength of different Fourier components with the model predictions. Such an approach may also be useful for higher resolution pupil phase retrieval schemes (47).

To map the optical transfer function, we project a series of SIM patterns with different frequencies and angles on a sample slide containing a thin layer of dye and observe the fluorescence at the camera (Fig. 1B). To avoid additional complications, we remove all polarization optics along the DMD path. For each pattern, we extract the amplitudes of the Fourier peaks at many reciprocal lattice vectors. We normalize the peak heights by the DC pattern component to correct for laser intensity drift. Finally, we compare the result to the Fourier components of the intensity pattern predicted by our forward model. The ratio of the measured and predicted peak value gives the optical transfer function of the imaging system. The results of this measurement are shown in Fig. 2.

**Fig. 1.**
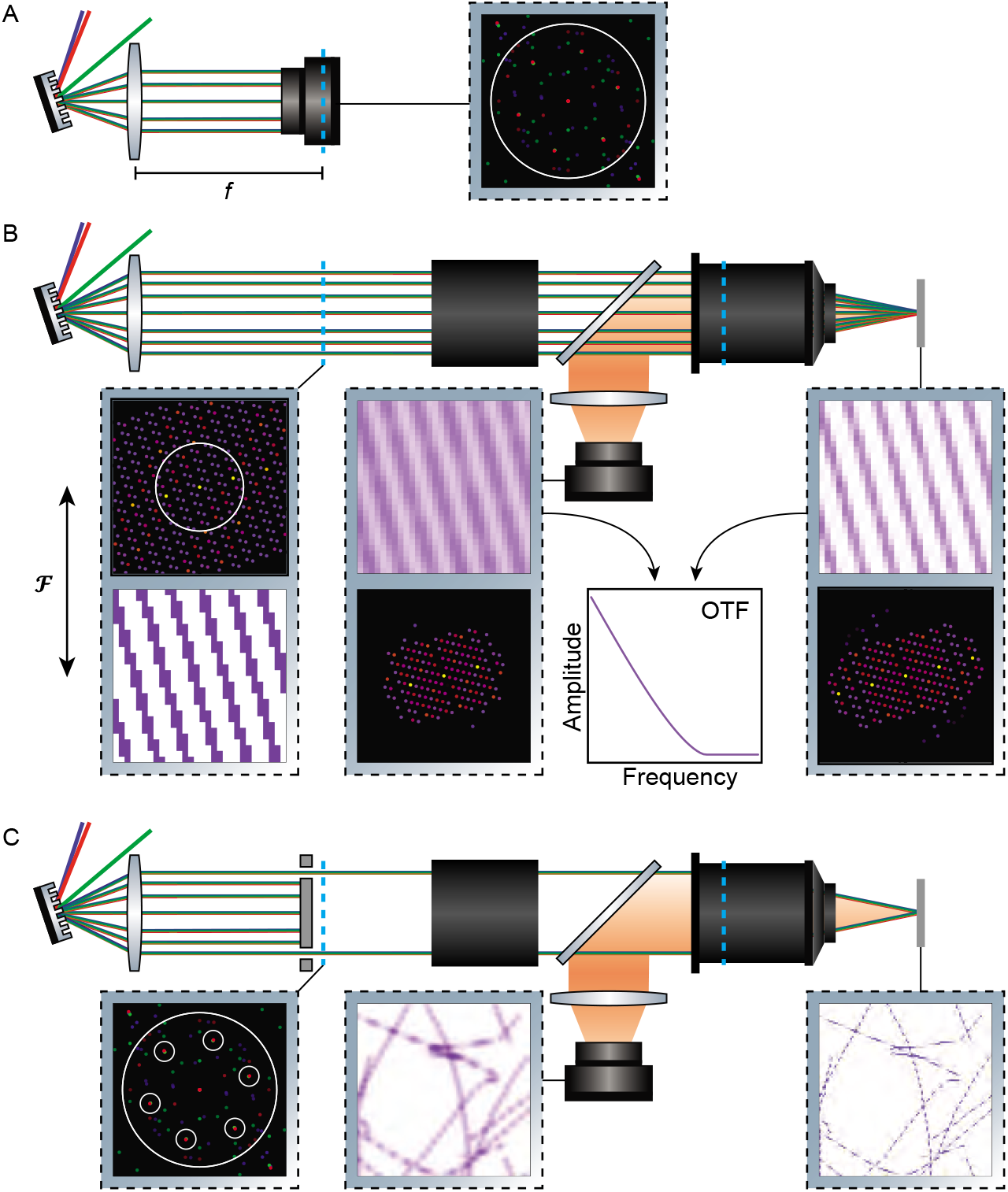
Experiment schematic. **A**. Simplified microscope layout for DMD and pattern forward model verification. The Fourier mask and polarization optics are removed from the system and the camera is placed in the back focal plane of the first lens, conjugate to the back focal plane of the objective (dashed blue line). **B**. Simplified microscope layout for OTF mapping. The Fourier mask and polarization optics are removed from the system, so all diffraction orders within a certain angular spread pass through the pupil (left inset). Dashed blue lines indicate the objective back focal plane and its conjugates. The diffraction orders are imaged onto a flat sample by additional optics (black box) and the objective, producing the pattern shown in the right inset. Fluorescence from the sample is imaged by the camera (center left inset). Predictions for the sample plane illumination based on the DMD and pattern projection forward models can be compared with the measured intensity in the camera plane to extract the system OTF (center right inset). In each inset, the top image shows the light intensity at a given point in the imaging system, while the bottom shows its Fourier transform. Forthe real space images low (high) intensity is shown in white (purple), while for the Fourier space images low (high) intensity is shown in black (colors increasing from purple to yellow). **C**. Simplified microscope layout when setup for SIM imaging. Three wavelengths of light diffract from the DMD and the diffraction is collected by the first imaging lens. A mask in the Fourier plane only passes the six diffraction orders required for projecting SIM patterns. The diffraction orders are shown for all nine SIM patterns (left inset) and 473nm (blue), 532 nm (green), and 635 nm (red) and the shape of the mask is illustrated by the white circles. Polarizers and additional optics are represented by the black box. The SIM patterns are projected onto the sample plane (right inset) and imaged through the microscope system producing a single raw frame (middle inset).

**Fig. 2.**
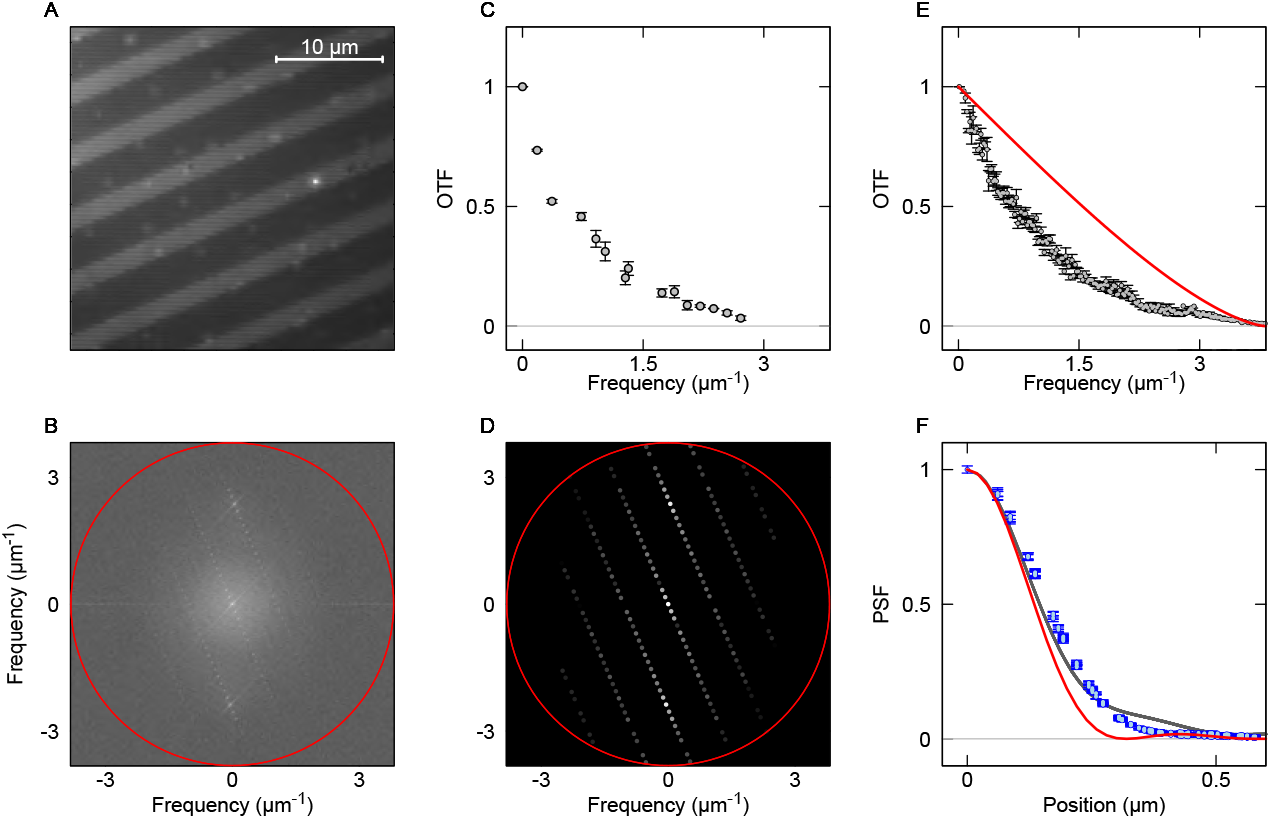
Experimental optical transfer function determination. **A**. Image of Alexa Fluor 647 dye slide excited with 473nm light and a DMD pattern with lattice vectors **r**_1_ = (−107, −106) and **r**_2_ = (111,117). **B**. Power spectrum of image from A. illustrating the discrete set of pattern diffraction frequencies. The red line denotes the maximum frequency passed by the imaging system. **C**. Peak heights obtained from the Fourier transform of A. divided by the expected intensity components of the DMD pattern. Only peaks that are expected to be larger than 1% of the DC value are shown. These provide an experimental estimate of the optical transfer function (gray points). Error bars are estimated from the noise power beyond the band cutoff. **D**. Theoretical power spectrum of the intensity pattern. The set of discrete frequency components predicted by the forward model matches that seen in the experiment. **E**. Experimental OTF (gray) determined with --3800 peaks from 360 patterns as in C. These points are binned and the error bars represent the standard error of the mean. The red line is the theoretical optical transfer function for a circular aperture and na = 1.3. **F**. Point spread functions corresponding to the OTF’s shown in E., including the ideal PSF for a circular aperture (red), the PSF obtained from the OTF (gray), and the PSF obtained from imaging diffraction limited beads (blue).

The optical transfer function can be estimated from

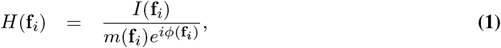

where *I* is the Fourier transform of the camera image, *H* is the optical transfer function, **f_*i*_** are the allowed Fourier components of the DMD pattern, and the *m* and *φ* are the amplitude and phase of intensity pattern generated by the DMD (Supplemental Note 4).

The experimental OTF rolls off more sharply than the ideal (Fig. 2E), which is expected for real optical systems. The point spread function obtained from the OTF agrees well with that obtained from diffraction limited beads (Fig. 2F). We use this experimental OTF in the reconstruction of our SIM data, which is expected to lead to more accurate reconstructions (Supplemental Note 9).

This approach can also be applied in real samples if additional corrections for sample structure are included. Incorporating this quantitative OTF measurement technique with adaptive optics would allow sensorless real-time aberration correction similar to what was achieved in (48), but incorporating additional information from our quantitative model.

While this approach is of general use, knowledge of the OTF is particularly important in SIM because the reconstruction algorithm is very sensitive to the optical transfer function, which is typically inferred from the point-spread function with low resolution because PSF’s of Nyquist sampled imaging systems have support on only a few pixels.

### 2.2. Estimating SIM resolution from variably spaced line pairs

We assessed the experimental SIM resolution by measuring the Argo-SIM test slide (Fig. 3). This slide includes test patterns consisting of variably spaced lines, ranging from 390 nm to 0 nm in steps of 30 nm. There are four of these patterns in different orientations (0°, 45°, 90°, 135°), allowing determination of the SIM resolution in all directions (Fig. 3A). We only assessed performance in the 473nm channel because the other channels do not efficiently excite fluorescence in the sample.

**Fig. 3.**
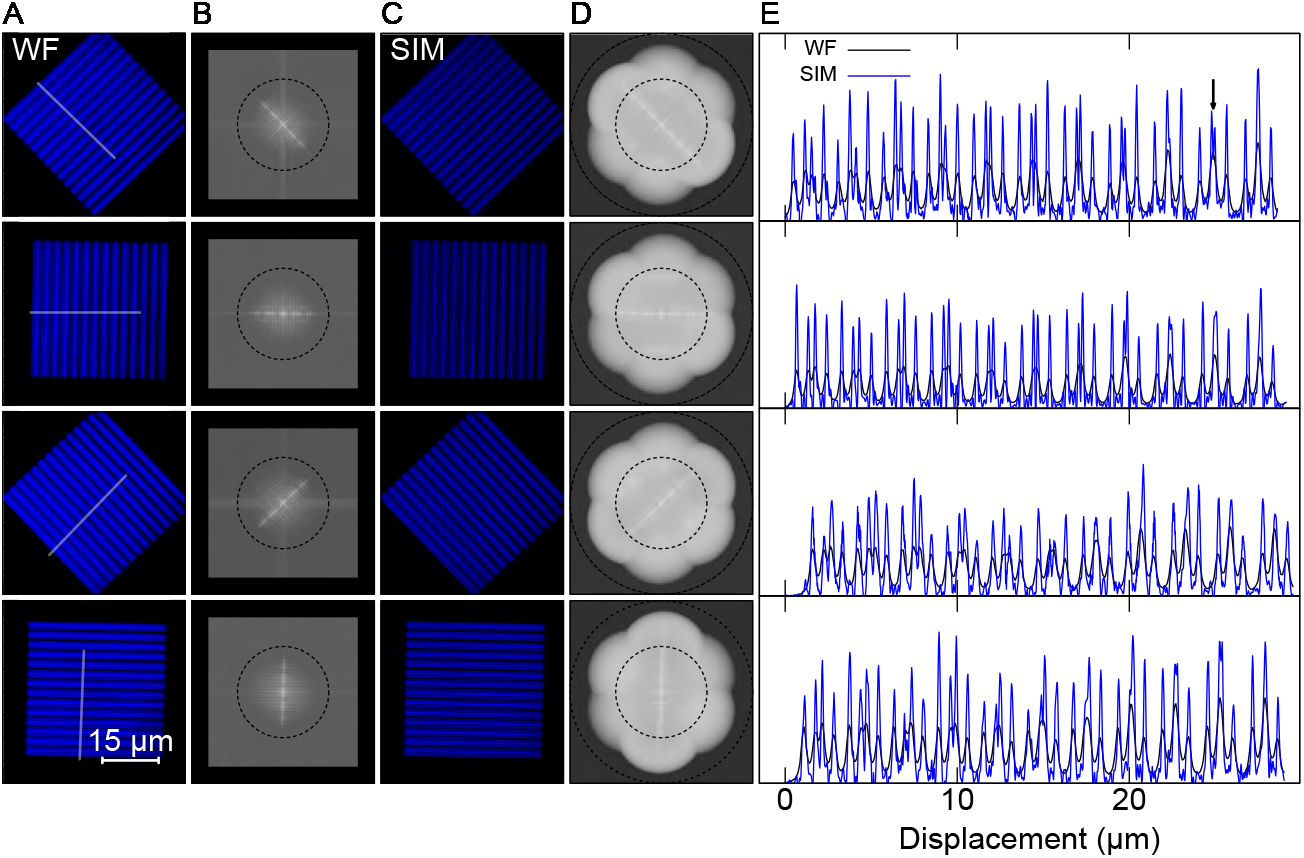
Experimental characterization of SIM resolution. **A**. Widefield images for 473 nm excitation light and different pattern orientations shown in analog-to-digital units (ADU). **B**. Power spectra of the images from A. The black circle illustrates the maximum frequency where the ideal optical transfer function has support for na = 1.3 and emission wavelength 519nm. For comparison with the SIM power spectrum, the widefield power spectrum is shown on the same frequency axes. Due to the sampling rate, the widefield image does not contain the highest frequencies present in the SIM image. We display a black background for these higher frequencies. **C**. SR-SIM images. **D**. Power spectra of SR-SIM images from D. The smaller circle as in B. The larger circle has twice the diameter, representing the maximum theoretical resolution for (linear) SIM. **E**. One-dimensional cuts plotted along the lines illustrated in A., showing 11 line pairs with spacings ranging from 390 nm to 90 nm in 30 nm steps. We show the widefield image (black) and SR-SIM image (blue). The 5th line pair, with spacing 270 nm, is the closest spaced pair visible in the widefield. The arrow in the top panel identifies 10th line pair, with spacing 120nm, which is the closest spaced line pair we distinguish in the SR-SIM reconstructions.

The smallest line pair we resolved is separated by 120 nm (10th pair, Fig. 3E). This should be compared with the minimum line spacing resolved by the widefield image, which is the 270 nm spaced pair (5th). However, the ability to resolve closely spaced objects is affected by contrast as well as resolution. To provide a quantitative and model-free characterization of SIM resolution, we assess the images using decorrelation analysis (49), which is available as an ImageJ plugin. Decorrelation analysis, which infers the resolution based on phase correlations in Fourier transform of the image, is expected to provide an accurate resolution estimate for sCMOS data, while Fourier ring correlation is not (50–52). For a conservative estimate of resolution enhancement, we report the relative resolution enhancement between the Wiener filtered widefield and SIM images. For the four images considered here the mean resolution enhancement is 1.54, and the maximum resolution enhancement is 1.61, corresponding to a resolution of ~130 nm. For an alternative test of instrument performance, we compared the size of diffraction limited fluorophores (see Supplemental Note 11.1), and found an enhancement of 1.71.

The experimental resolution should be compared with the upper theoretical bound on SIM resolution set by the SIM pattern spacing. The maximum theoretical resolution for *λ* ~ 520nm, na = 1.3, and SIM pattern period 250nm–260nm is ~110nm, or a factor of ~1.78 enhancement over widefield. The maximum possible resolution enhancement is reduced from 2 to ~1.78 because we use a SIM pattern frequency that is 71 % of the maximum value allowed for our objective na. The exact enhancement factor will depend on the Stokes shift for a given fluorophore. This choice of SIM pattern frequency is advantageous for reliable determination of the SIM pattern in realistic samples, where aberrations will tend to obscure a higher pattern frequency. The difference between the measured and theoretical resolution is due to the finite signal-to-noise ratio (SNR) of the experimental images. The highest frequency SR information falls at the edge of the OTF, and the intensity must exceed the noise to be detectable. At large spatial frequency, near the edge of the theoretical SIM resolution, the SNR drops below one and we attenuate both signal and noise from this region using a Wiener filter. This is illustrated in Fig. 3D, where the Wiener filter cuts off the Fourier space information at a lower frequency than the theoretical resolution (illustrated by the larger black circle). The non-ideal shape of the OTF causes a more rapid drop-off in signal strength than would naively be expected. In biological samples the achieved resolution is limited by the same effect, and hence is sensitive to available laser power, fluorophore brightness, sample aberrations, and SIM pattern modulation contrast. We further explored the role of signal-to-noise ratio and the use of our calibrated OTF via simulations (Supplemental Note 9). Although common practice, we do not apply post processing after SIM reconstruction (such as additional deconvolution or notch filtering) to enhance the reported SIM resolution.

### 2.3. Two-wavelength imaging of fixed cells

As an initial test of multicolor imaging using both “on” and “off” states of the DMD, we performed two wavelength SIM using the 473 nm and 532 nm channels to image actin filaments and mitochondria labeled with Alexa Fluor 488 and MitoTracker Red CMXRos in fixed bovine pulmonary artery endothelial (BPAE) cells. The SIM images substantially narrow the apparent width of the actin filaments (Fig. 4), in many cases making two filaments visible that were not distinguishable in the widefield image. The mitochondria similarly reveal cristae which cannot be distinguished in the widefield, but are visible in the SIM. Applying the decorrelation analysis, we find SIM leads to resolution enhancements by factors of ~1.5 and ~1.65 in the 473 nm and 532 nm channels respectively. We verified our reconstruction results using FairSIM (Supplemental Figure 4) and further explored the role of SNR in filamentous networks via simulation (Supplemental Note 9).

**Fig. 4.**
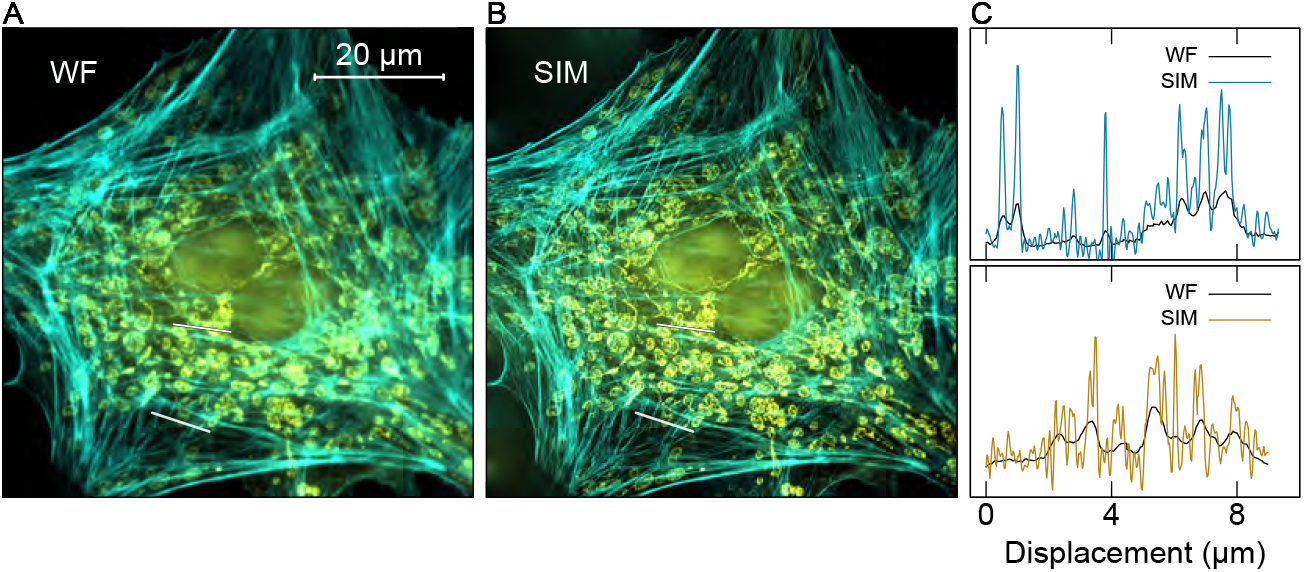
Two-color imaging of BPAE cells. **A**. Widefield image of a BPAE cell showing actin filaments in the 473 nm channel (cyan), and mitochondria in the 532 nm channel (yellow). Images are displayed in ADU. **B**. SIM-SR reconstruction corresponding to the image in A. **C**. One-dimensional cuts plotted along the lines illustrated in A. and B. We show the widefield image (black line) and SR-SIM image (colored line) for the 473 nm excitation (top) corresponding to the lower line in A. and 532 nm excitation (bottom) corresponding to the upper line in A. The SIM traces show significant enhancement of resolution and features which cannot be distinguished in the widefield image.

### 2.4. Three-wavelength imaging of live adenocarcinoma epithelial cells

We demonstrated time-resolved three-color SIM of live human adenocarcinoma cells by imaging mitochondria, actin, and lysosomes labeled with MitoTracker green, CellMask orange, and LysoTracker deep red. We imaged cell dynamics over a period of 15 min with a field of view of 100 μm × 90 μm, taking images at 1 min intervals (see Supplemental Movies 1 and 2). To increase SNR across the large FOV, we used a more powerful blue laser with nominal wavelength of 470nm instead of the 473nm laser used in the other SIM experiments (Supplemental Note 3). We chose an exposure time of 50 ms for raw SIM images, corresponding to 0.45 s per color and 1.65 s for a full three-color image. We chose the longest acquisition time such that mitochondria and lysosome dynamics were negligible during the 9 SIM images, which maximizes the SIM SNR. A single frame is shown in fig. 5, demonstrating enhanced contrast in the SIM image for the mitochondria and lysosomes and reveals branching of actin filaments which cannot be resolved in the widefield. Applying decorrelation analysis reveals resolution enhancement of 1.3,1.65, and 1.4 in the 470 nm, 532 nm, and 635 nm channels respectively. The blue channel has a significant amount of background from out-of-focus features. Specializing to the region shown in Fig. 5D where this is minimized, we find the resolution enhancement is somewhat higher, 1.37. The smaller resolution enhancement values reported here versus for the argo-SIM slide and fixed cells is likely a reflection of the lower SNR achieved in this measurement.

**Fig. 5.**
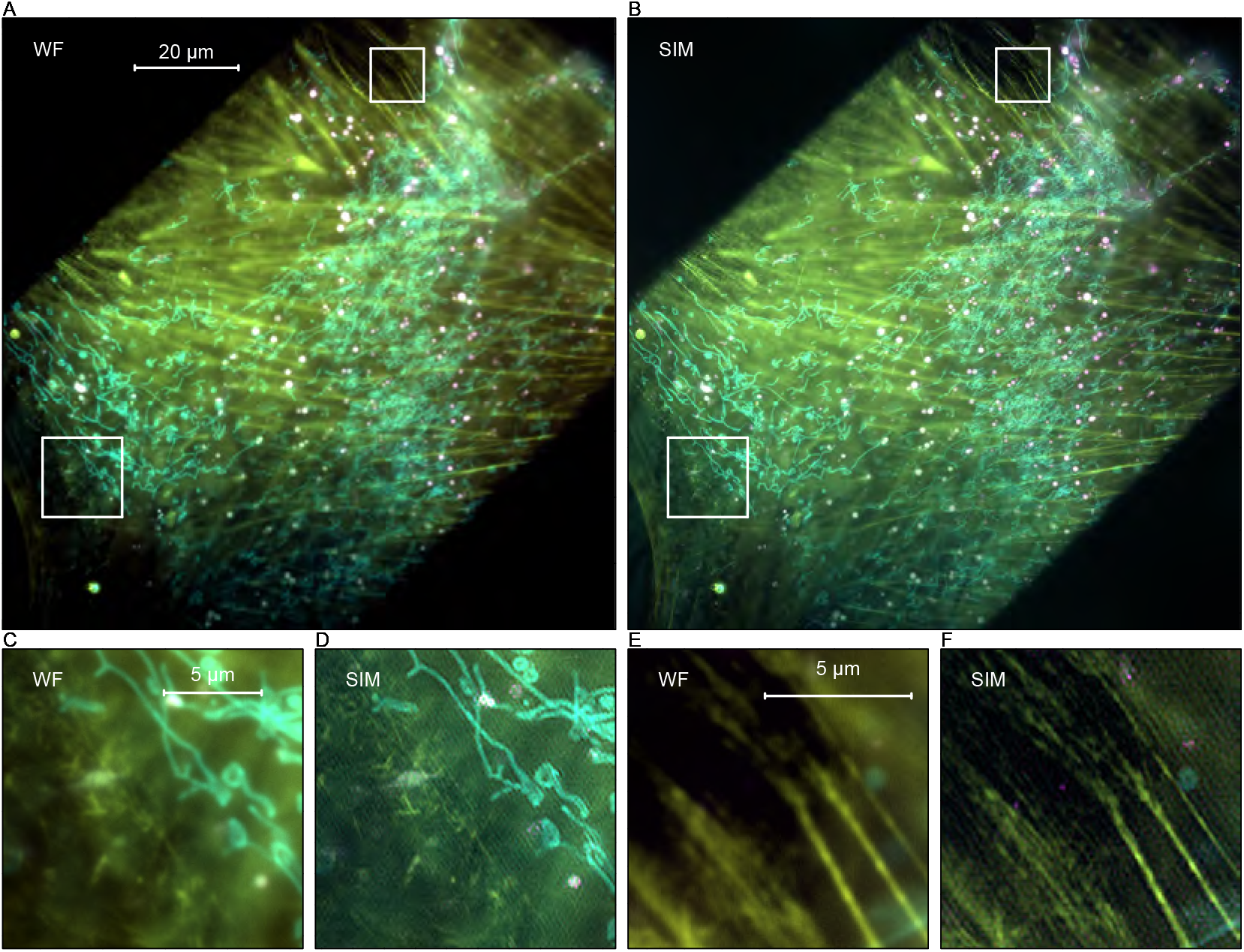
Three-color imaging of live human adenocarcinoma epithelial cells. **A**. Widefield image of human adenocarcinoma epithelial cell showing actin filaments (yellow) in the 532nm channel, lysosomes (purple) in the 635nm channel, and mitochondria (cyan) in the 470 nm channel. The field of view is limited by the size of the DMD, which is visible as the bright rectangular region in the image. Images are displayed in ADU. **B**. SIM-SR reconstruction corresponding to the image in A. **C., D**. Widefield and SIM images of the lower-left region of interest illustrating resolution enhancement for the mitochondria and actin filaments. Various short actin filaments that are difficult to see in the widefield image are visible in the SIM image with higher contrast, and their width is considerably narrowed. **E., F**. Widefield and SIM images of a the upper-right region of interest illustrating resolution enhancement for several longer actin filaments.

Our setup can realize a raw frame rate of ~330Hz, limited by the speed of the TriggerScope. Using a fast DAQ would allow us to reach the maximum exposure rate allowed by the DMD, 10 kHz, and replacing the DMD used here with the fastest available model could enable a 30 kHz frame rate. In most experiments sample properties such as SNR and fluorophore brightness impose significantly slower speed limits, but the high potential speed of DMD based setups could advantageous in some situations. For example, ferroelectric SLM based Hessian TIRF-SIM experiments have reached exposure times as low as 0.5 ms for a ~17 μm × 8 μm field of view, but slow SLM operation limited the raw frame rate to ~870 Hz (45).

## 3: Discussion

Multi-wavelength coherent SIM is regularly achieved using diffraction gratings or LCoS SLM’s for fringe projection, but the maximum achievable pattern display rate is limited by physical translation and rotation of the grating or the refresh rate of the SLM. Here, we provide a new theoretical framework to leverage a single DMD as a polychromatic diffractive optic for multi-wavelength coherent SIM, extending previous work (40,41). Our work significantly differs from previous SIM approaches using multi-wavelength incoherent LED light sources or multi-wavelength incoherent image projection using a DMD (42, 53). In the current work, the maximum resolution is governed by the coherent transfer function and does not require hundreds to thousands of raw images to generate a SIM image, or require scanning.

This opens the possibility for quantitative, multi-wavelength pattern formation at rates up to 30 kHz for a factor of ~5 lower cost than LCoS SLM based units. While SIM imaging rates are ultimately limited by signal-to-noise ratio and phototoxicity, fast control of multi-wavelength pattern formation also provides new avenues in the design of multi-wavelength tomography (1, 2), multi-wavelength optical trapping (3–5, 7), high-speed tracking of photostable fluorescent labels below the diffraction limit (54), and high-speed modulation enhanced localization microscopy in multiple colors (37, 43, 55).

Future improvements to our approach may include extension to more wavelengths (see Supplemental Note 1), 3D-SIM (35, 36), online GPU processing to speed reconstruction (34), specialized pattern generation to account for rolling shutters (56), and multiple cameras to speed multi-wavelength acquisition. Ease of alignment and instrument flexibility could be improved by removing the need for the Fourier mask. This could be achieved by developing a SIM reconstruction algorithm capable of accounting for the parasitic diffraction peaks, an approach that would require a detailed DMD forward model such as the one presented here. Implementing adaptive optics corrections could improveimaging quality and reduce artifacts, as has previously been reported (48, 57).

## 4: Materials and Methods

### 4.1. DMD diffraction forward model

To develop the DMD forward model, we calculate the diffracted light profile for an incident plane wave in the Fraunhofer approximation by considering the phase shift introduced by each point on the micromirror surfaces (40, 58–60). We adopt the same coordinate system as (40), where the DMD normal is along the 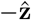 direction, the micromirrors are organized in a regular grid aligned with the 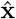 and 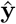 directions, and the micromirrors swivel about an axis along the 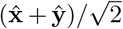 direction. For a plane wave of wave vector *k* = 2*π/λ* incident along direction **â**, the diffracted electric field in direction 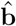 is

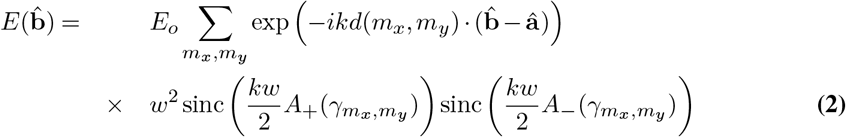

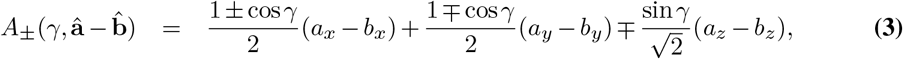

where *γ_m_χ_,m_y__* is the angle the mirror at position (*m_x_,m_y_*) makes with grating normal, *w* is the mirror width, and *d* is the spacing between adjacent mirrors. At each mirror *γ* takes one of two values: either 12° (“on”) or —12° (“off”). Here the sinc envelope expresses the effect of rays interfering from the same micromirror, and the sum represents rays interfering from different micromirrors.

To incorporate DMD diffraction into the overall optical system response, we recast the effect of the DMD as an effective pupil function. To do this we define the pattern function, *P(m_x_,m_y_*) where we take *P* = 0 (*P* = 1) at “off” (“on”) mirrors. We recognize that eq. 3 gives the discrete Fourier transform (DFT) of the pattern function, 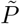, therefore the diffracted electric field becomes

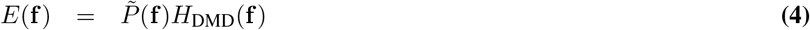

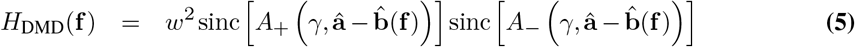

where **f** is the spatial frequency of the DMD image and 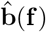 is the output unit vector diffracted by the pattern component at **f**. We ignore the “off” state mirrors in this equation, as we are only considering frequencies near a single solution of the diffraction condition, and the sinc envelopes for the “on” and “off” state mirrors are typically well separated.

The complete information about the pattern is contained in a discrete set of output angles (frequencies) given by the DFT frequencies

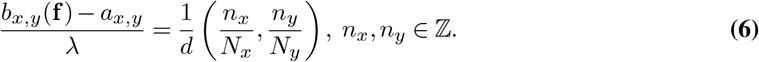

When *n_x_* (*n_y_*) is a multiple of *N_x_* (*N_y_*) eq. 6 is the diffraction condition for the underlying grating. Other frequencies are generated by diffraction from the DMD pattern. For a finite pixel grid, intermediate frequencies can be calculated from the DFT using an analog of the Whittaker-Shannon interpolation formula.

Equations 4–6 constitute a complete forward model of DMD diffraction. The expressions obtained here remove the need to numerically evaluate integrals or perform other expensive numerical simulations to determine DMD diffraction. It is only necessary to calculate a discrete Fourier transform, solve for output angles, and evaluate the sinc factor.

### 4.2. Multicolor DMD diffraction

We now apply our forward model to evaluate the constraints it places on multicolor operation. We first specialize to the plane in which the micromirrors swivel, i.e. the (*x — y*) *z* plane, which simplifies the analysis because light incident in this plane has its primary diffraction orders in the same plane. For light incoming and outgoing in this plane, the blaze and diffraction conditions reduce to (58),

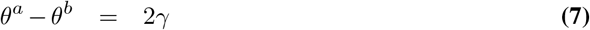

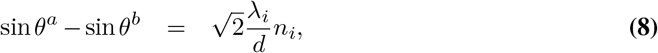

where *θ^a,b^* are the angles between **â** or 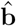 and the DMD normal in the (*x — y*) *z* plane and *i* indexes the different wavelengths. The blaze condition (eq. 7) is the angle where the law of reflection is satisfied for light incident on a single micromirror (58). For *N* wavelengths, this is a system of *N* + 1 equations with two angles and *N* diffraction orders as free parameters. In Supplemental Note 1 we show the blaze and diffraction conditions can be solved analytically for arbitrary input angles.

To realize multicolor operation we must solve this system for *N* > 1, which we achieve by first solving the blaze and diffraction conditions for *λ*_1_, and then attempting to satisfy 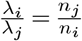. This ratio condition can be solved by finding rational approximations to each of these, 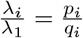. Then an approximate solution is obtained from *n*_1_ = lcm(*p*_1_ …,*p_N_*) and 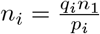. Any deviation between the rational approximation and the wavelength ratio must be accounted for by changing the input angle of the incident light, which entails slight violation of the blaze condition.

Additional colors can also be injected using the DMD mirror “off” state. Supposing we have already fixed the input and output angle for several colors using the “on” state mirrors, additional wavelengthsmust satisfy

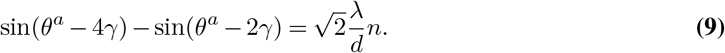

For our DMD parameters, *d* = 7.56μm and *γ* = 12°, two color operation can be achieved using *λ*_1_ = 473nm, *λ*_2_ = 635nm, by approximating *λ*_2_/*λ*_1_ ~ 4/3 which implies *n*_1_ = 4,*n*_2_ = 3, with *θ^a^* ~ 43° and *θ^b^* ~ 19°. Adding a third wavelength, *λ*_3_ = 532nm, using this approach is challenging because the smallest rational approximation with less than 10% error is *λ*_3_/*λ*_1_ ~ 8/7, implying *n*_1_ = 8, *n*_2_ = 7, *n*_3_ = 6. However, the maximum diffraction orders allowed by eqs. 7 and 8 are *n*_max_ = 4, 3, 3 respectively. Instead, we achieve three color operation by injecting 532 nm light using *θ^a^* = —3.5° and the —4th order from the “off” mirrors. Overlapping the 532 nm diffraction with the other colors requires a deviation of ~1.5° from the blaze condition. Perfect alignment of the —4th order occurs near 550 nm.

Deviations from the blaze condition degrade the SIM modulation contrast. To quantify this degradation, let *η* be the imbalance between the two components of the diffracted electric field that interfere to produce the SIM pattern. The pattern modulation contrast is the ratio of amplitudes of the high frequency and DC components,

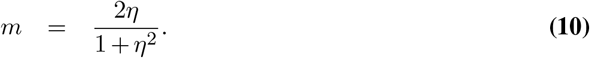

The contrast depends on the angle of the SIM pattern, ranging from a minimum along the *θ_x_* = —*θ_y_* direction to a maximum along the *θ_x_* = *θ_y_* direction. We summarize the worst case contrast for our parameters in table 1 and fig. 6, where we find high contrast is expected despite the modest violation of the blaze condition at 532 nm.

**Table 1.** Comparison of modulation contrast degradation in SIM patterns due to violation of the blaze condition versus wavelength for the 5 mirror period patterns shown in Fig. 6.

| wavelength | $\theta^a$ | diffraction order | $\eta^2$ | $m$ |
| --- | --- | --- | --- | --- |
| 473 nm | $43.66^\circ$ | (4, -4) | 1 | 1 |
| 532 nm | $-3.53^\circ$ | (-4, 4) | 0.66 | 0.979 |
| 635 nm | $43.86^\circ$ | (3, -3) | 0.95 | 0.9997 |

**Fig. 6.**
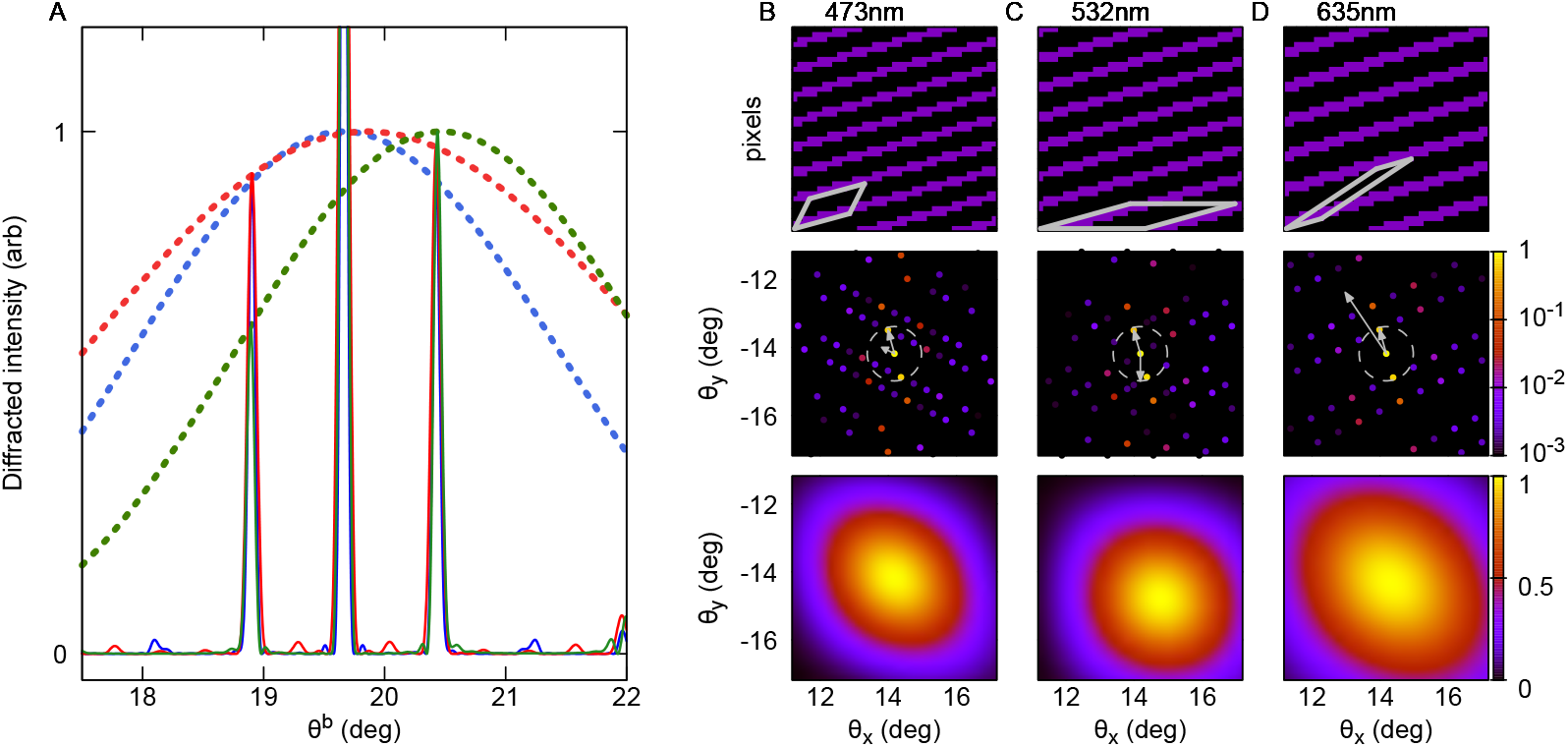
DMD diffraction forward model. **A**. Diffracted intensity for *θ^a^* = 43.66° and *λ* = 473nm (blue), *θ^a^* = −3.53° and 532nm (green), and *θ^a^* = 43.86° and 635 nm (red). For these input angles, 473 nm satisfies the Blaze condition, and 635 nm satisfies it to within a fraction of a degree. 532 nm shows a larger deviation, visible in the shift of the envelope center and the asymmetry in the diffraction peaks. To ensure the diffraction peaks for all colors appear at the same location and along the *θ_x_* = — *θ_y_* line the patterns simulated here have periods of 5, 5.62, and 6.71 mirrors which are not commensurate with the pixel lattice. **B**. DMD mirror pattern showing “on” mirrors (purple), “off” mirrors (black), and one unit cell (gray outline) for **r**_1_ = (11,3) and **r**_2_ = (3,6) (upper panel). Diffracted intensity from the DMD using 473nm light incident at *θ^a^* = 43.66° shown on a log scale. The maximum acceptance angle (gray circle) and reciprocal vectors (gray arrows) are shown (center panel). The envelope function illustrating where the blaze condition is satisfied, shown with a linear scale. **C**. panels the same as B, but for 532 nm light incident at —3.53° with **r**_1_ = (18,5) and **r**_2_ = (21,0). **D**. 635 nm light incident at 43.86° with **r**_1_ = (7, 2) and **r**_2_ = (18,12).

### 4.3. DMD SIM forward model

To quantitatively characterize SIM pattern formation in our system, we apply the DMD forward model to a set of mirror patterns commonly employed to generate SIM sinusoidal illumination profiles. SIM patterns are designed to be periodic to maximize diffraction into a single spatial frequency component. We define SIM patterns on a small subset of DMD mirrors, the unit cell, which is tiled across the DMD by a pair of lattice vectors, **r**_1_ and **r**_2_, which have integer components. This produces a periodic pattern in the sense that mirrors separated by *n***r**_1_ + *m***r**_2_ for any 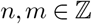 will be in the same state.

The lattice structure implies all frequency components of the SIM pattern are multiples of the reciprocal lattice vectors, **k**_1,2_, defined by the property **r**_*i*_ · **k**_*j*_ = *δ_ij_* (61). This constrains the frequencies of the diffracted electric field to the set **f** = *n*_1_**k**_1_ + *n*_2_**k**_2_. Furthermore, the Fourier components can be calculated from the unit cell *U*, which is typically much smaller than the full DMD,

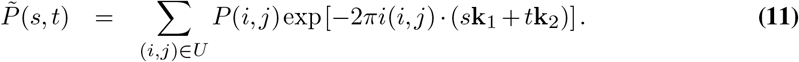

This expression is correct up to a boundary term when the unit cell does not perfectly tile the DMD. To generate appropriate SIM patterns for the experiment we construct **r**_1,2_ such that **k**_2_ matches a desired period *P* and angle *θ* (see Supplemental Note 6). We also require that the pattern can be translated to change its phase, which is achieved by setting **r**_2_ = (*n, m*)*n_p_* where *n_p_* is the number of desired SIM phases (31, 32, 35, 36). Next, we construct one unit cell of our pattern by generating a smaller cell *U_p_* from the vectors **r**_1_ and **r**_2_/*n_p_*. By construction *U_p_* is contained in *U*. Setting all pixels in *U_p_* to “on” and all other pixels in *U* to “off” creates the desired pattern, which has Fourier components

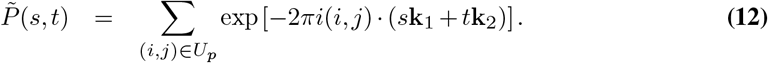

The strongest Fourier component of this pattern occurs at **k**_2_, which defines the SIM frequency. However the pattern also has Fourier weight at other reciprocal lattice vectors due to its binary and pixelated nature. These additional Fourier components introduce unwanted structure in the SIM patterns, and are blocked in the experiment by inserting a mask in the Fourier plane.

Although these Fourier components are unwanted in SIM operation, our ability to precisely predict their amplitude and frequency can be used to extract information about either our optical system or a sample of interest. These predictions are highly accurate (Supplemental Note 2), and hence can be used to map out the frequency response of our imaging system. In the future this information could also be harnessed to enhance SIM reconstruction.

### 4.4. SIM instrument design

One objective of our design was to enable ease of use and to minimize effort needed adopt, modify, and develop new imaging experiments using multiple coherent light sources and a DMD. With this in mind, our multicolor SIM system was assembled entirely from commercially available parts, easily machined or 3D printed parts, and standard optical components. Coherent light was supplied by diode and diode-pumped solid-state (DPSS) lasers made by Lasever. We utilized a blue laser (either 470 nm or 473 nm), green laser (532 nm), and red laser (635 nm). We used the 473nm laser for the argo-SIM slide resolution test and two-color fixed cell imaging. We used the 470 nm laser for the three-color live-cell imaging, OTF measurement, and DMD model validation. Although the nominal wavelength of the OEM 470nm-2W laser is 470 nm, we estimate its actual wavelength is ~465 nm based on the transmitted fraction of light from a dichroic mirror with sharp edge in the blue. This wavelength is confirmed by the model fit in Supplemental Note 2. The various colors were combined with dichroic mirrors (M1-8, DM1, DM2) and coupled into a square-core multimode optical fiber with core size 150 μm (MMF) using a reflective coupler (FC1). The fiber was shaken using an agitator module (62) to reduce the influence of speckle.

The output from the fiber (FC2) was imaged onto the surface of a dual-axis voice-coil mirror (VCM) using an *f* = 30mm achromat (L1). A voice-coil mirror was used because of the large mirror area (15 mm) compared to standard available galvonometer mirrors. The surface of the voice-coil mirror was imaged onto a DMD using a scan lens (L2) and an *f* = 400 mm achromat (L3). The DMD was imaged onto the sample plane using a series of two 4*f* imaging systems. The first imaging system was formed by a tube lens (L6) and an *f* = 400 mm achromat (L7). After passing through the tube lens, the various SIM orders were linearly polarized in the azimuthal direction using a segmented (“pizza”) polarizer placed in the Fourier plane (POL), and unwanted diffraction orders were removed with a home built aluminum foil Fourier mask mounted on a lens tube (FM). The alignment of both is critical, and so we mounted them on XY-stages (Thorlabs, CXYZ-1 and Thorlabs, CXY2). The second imaging system was formed by an *f* = 300 mm achromat (L8) and the objective lens (OBJ). Before passing through the objective, the incoming light was filtered and reflected off a dichroic mirror (DM4). The emitted fluorescence light passed through the dichroic mirror and additional filters (DM4, EmF) and was imaged onto the camera (CAM) using a tube lens (L9). To improve SIM interference contrast, we use two matched dichroic mirrors (DM3, DM4) placed at a right angle to each other in the system. This is intended to counteract the differential phase shift of a single dichroic on the *s-* and *p*-polarization components of the SIM excitation light (32). The microscope body was an Olympus IX71 with the factory excitation paths and sample stages removed. 3D sample manipulation was achieved using a motorized XY-stage (Sample stage) and a piezo Z-stage (Objective piezo). A complete diagram of the apparatus can be found in Supplemental Note 5.

Our goal for ease of use extends to the control and reconstruction software. We created a control scheme that runs underneath the open-source project Micromanager 2.0 Gamma (63) and used a low-cost digital triggering device (Advanced Research Consulting Triggerscope 3B). The experiment was controlled using a desktop computer (Lenovo, ThinkServer TS140) running Microsoft Windows 10 Pro with the software Micro Manager 2.0 Gamma through the BeanShell scripting interface. Users set up multi-dimensional acquisitions as normal in Micromanager 2.0 Gamma and then ran our custom script to execute SIM imaging. This custom script bypassed the standard Micromanager acquisition engine. Instead, we only utilized the Micromanager ring buffer to acquire frames from the camera and the rest of the acquisition is hardware controlled. TTL triggering of the DMD and lasers as well as analog control of the voice coil mirror and objective piezo was accomplished with microsecond timing using the Arduino Due based Triggerscope with custom firmware. The DMD was controlled over USB using custom Python software. The SIM patterns were preloaded on the DMD firmware using the Texas Instruments DLP LightCrafter 6500 and 9000 GUI 4.0.1. We hope future development in this area continues to push towards making high-speed, deterministic hardware triggered microscopy widely available (64).

All instrument control code is available on GitHub https://github.com/QI2/mcSlM.

**Table 2.** Optical elements

| Component | Supplier | Part number |
| --- | --- | --- |
| Blue laser | Lasever | LSR473NL-150 |
| Blue laser | Lasever | OEM 470nm-2W |
| Green laser | Lasever | LSR532NL-500 |
| Green laser | Lasever | LSR532H-2W |
| Red laser | Lasever | LSR635-500 |
| M1-M13 | Thorlabs | BB1-E02 / BB2-E02 |
| DM1 | Semrock | LM01-552-25 |
| ExF | Semrock | LF405-488-532-635-B-OMF |
| DM2 | Semrock | LM01-480-25 |
| FC1 | Thorlabs | RC02FC-P01 |
| MMF | Thorlabs | M103L05 |
| Fiber shaker | Custom | Ref (62) |
| FC2 | Thorlabs | SM1FC |
| L1 | Thorlabs | AC254-30-A-ML |
| VCM | Optotune | MR-15-30 |
| L2 | Thorlabs | CLS-SL |
| L3 | Thorlabs | AC508-400-A-ML |
| L4 | Thorlabs | AC254-075-AB-ML |
| L5 | Thorlabs | AC508-500-A-ML |
| DMD | Texas Instruments | DLP Light Crafter 6500, DLP6500FYE |
| L6 | Nikon | MXA20696 |
| FM | Custom | — |
| POL | Codixx | colorPol vis 500 BC4 CW01 |
| DM3 | Chroma | zt473/532/633rdc-uf3 |
| L7 | Thorlabs | AC508-400-A-ML |
| L8 | Thorlabs | AC508-300-A-ML |
| DM4 | Chroma | zt473/532/633rdc-uf3 |
| EmF | Semrock | LF405-488-532-635-B-OMF |
| EmF | Chroma | ZET457NF |
| OBJ | Olympus | UPlanFL N 100x, $n_a = 1.3$ |
| Objective piezo | Mad City Labs | Nano F-200S |
| Sample stage | Mad City Labs | MicroDrive |
| L9 | Olympus | SWLTU-C |
| CAM | Hamamatsu | Orca Flash4.0 v2 C11440 |

### 4.5. Commercial samples for instrument characterization

Fluorescent DNA origami with localized fluorophores separated by 120 nm were used to quantify the initial alignment, point spread function (PSF), and SIM performance (Gattaquant DNA Nanotechnologies STED 120R). Additional calibration was done during alignment using multicolor fluorescent microspheres (Thermo Fisher T14792). A fluorescent slide with patterns intended to evaluate instrument resolution was used to characterized and fine tune SIM performance (Argolight Argo-SIM). A commercial slide with bovine pulmonary arotic endothelial (BPAE) cells labeled for nuclei (DAPI), *β*-actin (Phalloidin – Alexa488), and mitochondria (MitoTracker Red CMXRos) was used to evaluate and fine tune initial multiple wavelength performance (Thermo Fisher F36924).

### 4.6. Live adenocarcinoma epithelial cells

MDA-MB-231, human breast adenocarcinoma, cells (ATCC HTB-26) were cultured on 50 mm diameter #1.5 glass bottomed cell culture dishes (World Precision Instrument, FD5040100) at 37 °C in L-15 Medium (ATCC, 30-2008) with 10 % FBS (ATCC, 30-2020) and 5 % CO2 for a minimum of 24-36 hours to allow for adherence and acclimation. Live cells were then stained for 30 min at 37 °C in 5mL L-15 medium + 10% FBS with 100 nM MitoTracker Green FM, 2x CellMask Actin Orange Tracking Stain (Invitrogen, 2212429) and 100 nM LysoTracker Deep Red (Invitrogen, L12492). Stain and medium were aspirated off and replaced with 1 mL-2 mL fresh L-15 + 10 % FBS medium. Stained cells were maintained for a short period of time in a 37 °C incubator until imaging.

## Supporting information

Supplemental Movie 1 (widefield)

Supplemental Movie 2 (SIM)

## Acknowledgments

We thank Drs. Randy Bartels, Thomas Huser, Marcel Müller, and Andrew York for helpful discussions on the use of a DMD as an active diffractive element.

## Author contributions

Theoretical model, numerical simulations, hardware design and construction, software development, and instrument validation: PTB. Sample preparation and imaging: RK, GJS, PTB, DPS. Original concept, project management, and writing of manuscript: PTB, DPS.

