## Supplemental Information for "Multicolor structured illumination microscopy and quantitative control of polychromatic coherent light with a digital micromirror device"

### Contents

**1 General closed-form solution of the blaze and diffraction conditions** **1**

**2 DMD model validation** **4**

**3 DMD diffraction efficiency** **6**

**4 OTF measurement technique** **7**

**5 SIM instrument design** **8**

**6 Multicolor pattern generation algorithm** **10**

**7 SIM reconstruction** **11**

**8 Comparison of SIM reconstruction with FairSIM** **12**

**9 SIM reconstruction of synthetic data** **13**

### 1: General closed-form solution of the blaze and diffraction conditions

Here we present a closed form solution to the combined blaze and diffraction problem, which removes any need to perform numerical simulations to determine appropriate input/output angles and diffraction orders for determining optimal points for multicolor operation. Further, it allows us to answer various useful questions which have not previously been addressed, including which diffraction orders can be overlapped with the blaze condition in principle.

To find a joint solution to the blaze and diffraction conditions, it is convenient to work directly with the incoming and outgoing unit vectors $\hat{a}$ and $\hat{b}$ as opposed to parameterizing these by angles, which corresponds to working with the direction cosines (1). However, for later convenience we also introduce

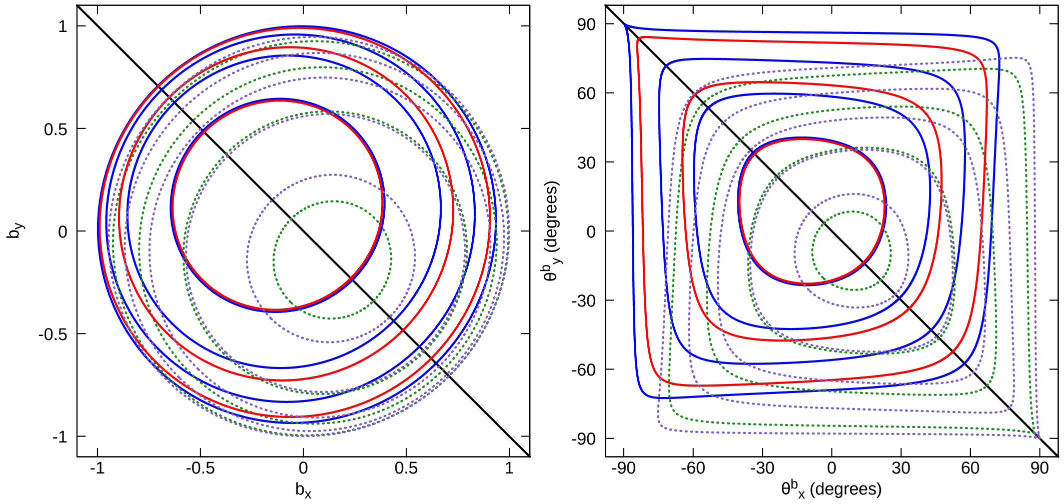

**Fig. 1. Blaze and diffraction condition solution contours. A**. Output diffraction angles parameterized by the components of $\hat{\mathbf{b}}$ for "on" mirrors and $473\,\mathrm{nm}$ for $n = 1, ..., 4$ (blue) and $635\,\mathrm{nm}$ for $n = 1, ..., 3$ (red), and solutions for "off" mirrors and $532\,\mathrm{nm}$ for $n = -4, ..., -1$ (green) and $405\,\mathrm{nm}$ for $n = -5, ..., -1$ (purple) in terms of the unit vector components $b_x$ and $b_y$. The shapes with smaller radii correspond to the higher diffraction orders. The black line indicates the $\theta_x = -\theta_y$ direction. **B**. Diffraction output angle data from A. parameterized by $\theta_x$ and $\theta_y$.

an angular parameterization following (2),

$$\hat{\mathbf{a}} = \left(\tan\theta^a_n \hat{\mathbf{n}} + \tan\theta^a_{n_\perp} \hat{\mathbf{n}}_\perp + \hat{\mathbf{z}}\right) / \sqrt{\tan^2\theta^a_n + \tan^2\theta^a_{n_\perp} + 1} \quad (1)$$

$$\hat{\mathbf{b}} = \left(\tan\theta^b_n \hat{\mathbf{n}} + \tan\theta^b_{n_\perp} \hat{\mathbf{n}}_\perp - \hat{\mathbf{z}}\right) / \sqrt{\tan^2\theta^b_n + \tan^2\theta^b_{n_\perp} + 1} \quad (2)$$

where we most often consider the bases $\hat{\mathbf{n}}, \hat{\mathbf{n}}_\perp = \hat{\mathbf{x}}, \hat{\mathbf{y}}$ and $\hat{\mathbf{n}}, \hat{\mathbf{n}}_\perp = (\hat{\mathbf{x}} - \hat{\mathbf{y}})/\sqrt{2}, (\hat{\mathbf{x}} + \hat{\mathbf{y}})/\sqrt{2}$. $\theta_n$ is the angle between the $\hat{\mathbf{n}}$ axis and the unit vector $\hat{\mathbf{a}}$ or $\hat{\mathbf{b}}$ projected onto the $\hat{\mathbf{n}}_\perp - \hat{\mathbf{z}}$ plane.
Now, we exploit the fact that the blaze condition is synonymous with the law of reflection being satisfied on a single micromirror. To more easily evaluate this condition, we adopt the basis

$$\hat{\mathbf{e}}_1 = \frac{1}{\sqrt{2}}(\hat{\mathbf{x}} - \hat{\mathbf{y}})\cos\gamma - \hat{\mathbf{z}}\sin\gamma \quad (3)$$

$$\hat{\mathbf{e}}_2 = \frac{1}{\sqrt{2}}(\hat{\mathbf{x}} + \hat{\mathbf{y}}) \quad (4)$$

$$\hat{\mathbf{e}}_3 = \frac{1}{\sqrt{2}}(\hat{\mathbf{x}} - \hat{\mathbf{y}})\sin\gamma + \hat{\mathbf{z}}\cos\gamma, \quad (5)$$

which is chosen such that $\hat{\mathbf{e}}_3$ is the micromirror normal. In this basis, the blaze condition is

$$a_1 - b_1 = 0 \quad (6)$$

$$a_2 - b_2 = 0 \quad (7)$$

$$a_3 + b_3 = 0., \quad (8)$$

which is equivalent to $A_\pm(\gamma, \hat{\mathbf{a}} - \hat{\mathbf{b}}) = 0$.
The general diffraction condition occurs when the exponentials in eq. 3 of the main text all enter with the same phase, and can be written

$$a_x - b_x = \frac{\lambda}{d} n_x \quad (9)$$

$$a_y - b_y = \frac{\lambda}{d} n_y, \quad (10)$$

which in the new basis becomes

$$(a_1 - b_1)\frac{\cos\gamma}{\sqrt{2}} + (a_2 - b_2)\frac{1}{\sqrt{2}} + (a_3 - b_3)\frac{\sin\gamma}{\sqrt{2}} = \frac{\lambda}{d} n_x \quad (11)$$

$$-(a_1 - b_1)\frac{\cos\gamma}{\sqrt{2}} + (a_2 - b_2)\frac{1}{\sqrt{2}} - (a_3 - b_3)\frac{\sin\gamma}{\sqrt{2}} = \frac{\lambda}{d} n_y. \quad (12)$$

Adding and subtracting these two equations, we find

$$a_2 - b_2 = \frac{\lambda}{\sqrt{2}d}(n_x + n_y) \tag{13}$$

$$(a_2 - b_2)\cos\gamma + (a_3 - b_3)\sin\gamma = \frac{\lambda}{\sqrt{2}d}(n_x - n_y) \tag{14}$$

Inserting eqs. 6–8, we find that the blaze condition and diffraction conditions can only be satisfied together for $n = n_x = -n_y$, and in this case

$$a_3 = \frac{1}{\sqrt{2}\sin\gamma}\frac{\lambda}{d}n \tag{15}$$

$$a_2 = \pm\sqrt{1 - a_3^2 - a_1^2} \tag{16}$$

$$a_1 \in \left[-\sqrt{1-a_3^2}, \sqrt{1-a_3^2}\right], \tag{17}$$

and two numbers, $n$ and $a_1$ completely specify the solution. Further the condition $0 < a_3 \leq 1$ implies this equation can only be satisfied for a finite number of $n$,

$$n = \begin{cases} 1, \ldots, \lfloor \frac{d}{\lambda}\sqrt{2}\sin\gamma \rfloor & \text{if } \gamma < 0 \\ \lceil -\frac{d}{\lambda}\sqrt{2}\sin\gamma \rceil, \ldots, -1 & \text{if } \gamma > 0, \end{cases} \tag{18}$$

where $\lfloor \cdot \rfloor$ and $\lceil \cdot \rceil$ are the floor and ceiling functions respectively.
For any solution, we can convert eqs. 15–17 back to the $\hat{\mathbf{x}}$, $\hat{\mathbf{y}}$, $\hat{\mathbf{z}}$ basis using eqs. 3–5. We explicitly perform this substitution for the case of most interest, $a_x = -a_y$ ($\theta_x = -\theta_y$), which implies $a_2 = 0$. Therefore,

$$\hat{\mathbf{a}} = \frac{\hat{\mathbf{x}} - \hat{\mathbf{y}}}{\sqrt{2}}\sin(\alpha + \gamma) + \hat{\mathbf{z}}\cos(\alpha + \gamma) \tag{19}$$

$$\hat{\mathbf{b}} = \frac{\hat{\mathbf{x}} - \hat{\mathbf{y}}}{\sqrt{2}}\sin(\alpha - \gamma) - \hat{\mathbf{z}}\cos(\alpha - \gamma) \tag{20}$$

where

$$\cos\alpha = \frac{1}{\sqrt{2}\sin\gamma}\frac{\lambda}{d}n \tag{21}$$

or, writing in terms of the parameterization eq. 1–2 for $(\hat{\mathbf{x}} - \hat{\mathbf{y}})/\sqrt{2}$, $(\hat{\mathbf{x}} + \hat{\mathbf{y}})/\sqrt{2}$, $\hat{\mathbf{z}}$,

$$\theta^a_{x-y} = \alpha + \gamma \tag{22}$$

$$\theta^b_{x-y} = \alpha - \gamma, \tag{23}$$

which gives *two* explicit solutions for the input and output angles in terms of only $n$ and $\gamma$, one for each solution to eq. 21.
For multicolor operation, we must solve eqs. 15–17 for multiple wavelengths, $\lambda_1, \ldots, \lambda_n$ subject to the condition $\hat{\mathbf{b}}_1 = \hat{\mathbf{b}}_2 = \ldots = \hat{\mathbf{b}}_n$. We see from these equations that for fixed value of $\gamma$ this requires $\lambda_1 n_1 = \ldots = \lambda_n n_n$, which can be solved using the algorithm described in the main text. The only continuously tuneable parameters here are the wavelengths, which is undesirable because wavelength choices will often be dictated by choice of fluorophore, existing equipment, or available laser sources. On the other hand, we can generally find a solution for two colors using the "on" and "off" mirror states of the DMD. In this case, let the wavelength and diffraction order for the "on" ("off") mirrors be $\lambda_p$ and $n_p$ ($\lambda_m$ and $n_m$). Suppose $\gamma = \gamma_{\text{on}} = -\gamma_{\text{off}}$. Then $\hat{\mathbf{b}}_p = \hat{\mathbf{b}}_m$ gives the quadratic equation

$$b_z = \frac{1}{2\sqrt{2}\sin\gamma\cos\gamma}\frac{-\lambda_p n_p + \lambda_m n_m}{d} \tag{24}$$

$$b_x^2 + b_x\left(\frac{\lambda_p n_p + \lambda_m n_m}{2\sin^2\gamma d}\right) = -\frac{1}{2}\left[b_z^2 + \left(\frac{\lambda_p n_p + \lambda_n n_m}{2\sin^2\gamma d}\right)^2 - 1\right] \tag{25}$$

which can be satisfied by tuning the input angles.

To develop a three-color solution for $473\,\mathrm{nm}$, $635\,\mathrm{nm}$, and $532\,\mathrm{nm}$, we apply both approaches described above. First, we take advantage of $473 \times 4 \approx 635 \times 3$ to nearly overlap these diffraction orders. Next, we take advantage of the $n = -4$ diffraction order of the $532\,\mathrm{nm}$ light using the "off" state mirrors. Eqs. 24 and 25 show the $473\,\mathrm{nm}$ and $532\,\mathrm{nm}$ outputs overlap at the out-of-plane position $(b_x, b_y) = (0.381, 0.0198)$, $(\theta^b_x, \theta^b_y) \sim (22.39°, \sim 1.22°)$, or $(\theta^b_{x-y}, \theta^b_{x+y}) \sim (17.04°, 15.44°)$. A similar observation has been made in (3), but without the benefit of a closed form solution.

Four-color operation can be realized using a similar approach, taking advantage of the near overlap between the $405\,\mathrm{nm}$ $n = -4$ and $532\,\mathrm{nm}$ $n = -3$ diffraction orders and keeping the $473\,\mathrm{nm}$ $n = 4$ and $635\,\mathrm{nm}$ $n = 3$ diffraction orders. Here, solving again for the overlap of the $532\,\mathrm{nm}$ and $473\,\mathrm{nm}$ beams, we find $(b_x, b_y) = (0.130, 0.583)$, $(\theta^b_x, \theta^b_y) \sim (9.21°, \sim 36.00°)$, or $(\theta^b_{x-y}, \theta^b_{x+y}) \sim (-21.76°, 32.15°)$. We illustrate this solution and the set of all joint solutions of the blaze and diffraction conditions for $405\,\mathrm{nm}$, $473\,\mathrm{nm}$, $532\,\mathrm{nm}$, and $635\,\mathrm{nm}$ in Fig. 1.

These solutions are promising for improved 3- and 4-color operation, but they require a substantially more complicated mounting structure for the DMD. Because the deviation from the blaze condition for our chosen 3-color configuration leads to negligible degradation of the SIM pattern, we believe it is a superior compromise. For achieving 4-color operation, working with a more complicated experimental configuration may be unavoidable. However, using more extreme angles poses other problems in the optical setup, in particular the tilt of the DMD may exceed the depth of field of the first imaging lens.

### 2: DMD model validation

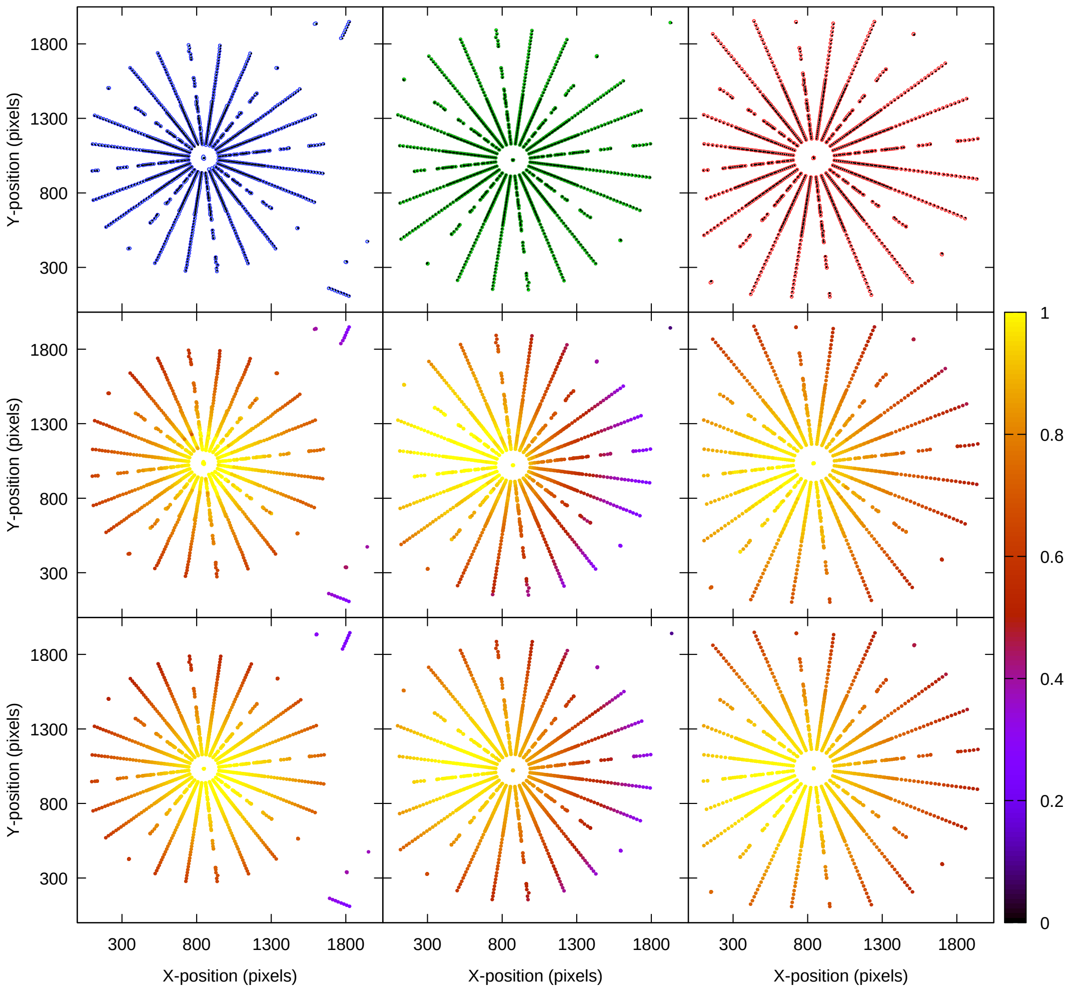

**Fig. 2. Model validation** Comparison between Fourier peaks measured in the imaging system pupil and the DMD and SIM pattern forward models. **A**. Comparison of measured positions (colored circles) and theoretical positions (black symbols) for nominal wavelengths $470\,\mathrm{nm}$ (left), $532\,\mathrm{nm}$ (middle), and $635\,\mathrm{nm}$ (right). **B**. Measured intensity values across the pupil normalized by the DC value and corrected for the expected diffraction intensity for each frequency component. **C**. Theoretical intensity values across the pupil.

| quantity | fit value | expected value |
|---|---|---|
| $\lambda_1$ | 464.93 nm | 465 nm |
| $\hat{\mathbf{b}}_1$ | (0.2524, -0.2584, -0.9325) | (0.2557, -0.2557, -0.9323) |
| $\lambda_2$ | 532.70 nm | 532 nm |
| $\hat{\mathbf{b}}_2$ | (0.2522, -0.2576, -0.9328) | (0.2557, -0.2557, -0.9323) |
| $\lambda_3$ | 635.84 nm | 635 nm |
| $\hat{\mathbf{b}}_3$ | (0.2525, -0.2587, -0.9324) | (0.2557, -0.2557, -0.9323) |
| $M$ | 31 001 | 30 769 |
| $\theta_c$ | 50.05° | — |
| $\hat{\mathbf{p}}$ | (0.2524, -0.2582, -0.9325) | (0.2557, -0.2557, -0.9323) |
| $(c_x, c_y)$ | (853.9, 1029.4) | — |
| $\theta_n$ | 44.47° | 45° |
| $d_x$ | 7.598 µm | 7.56 µm |
| $d_y$ | 7.558 µm | 7.56 µm |
| $w_x$ | 7.598 µm | — |
| $w_y$ | 7.558 µm | — |
| $\gamma_{\text{on}}$ | 12.05° | 12° |
| $\gamma_{\text{off}}$ | $-11.96°$ | $-12°$ |

**Table 1.** DMD parameters determined by non-linear least squares fit. Although the nominal wavelength of the blue laser used here is 470 nm, we have verified independently that its actual wavelength is ∼465 nm. See the materials and methods section for further discussion.

To validate the DMD and SIM pattern formation forward models presented in eqs. 3, 6, and 12 of the main text we measure the light diffracted by the DMD for 360 SIM patterns constructed as described in the materials and methods section of the main text and supplemental note 6, and compare the resulting diffraction order positions and intensities to our model. To accomplish this, we place a sCMOS camera in the pupil plane following lens L6 (see Supplemental Note 5) and record the intensities of a number of diffraction peaks for the various DMD patterns using three wavelengths, 470 nm, 532 nm, and 635 nm. We parameterize these peaks by the DMD Fourier frequency, $\mathbf{f}$, which generates them and perform Gaussian fits to the resulting image to determine their positions, $\mathbf{r}^e_c(\mathbf{f}) = (x^e_c(\mathbf{f}), y^e_c(\mathbf{f}))$ and intensities, $I^e(\mathbf{f})$ on the camera. We compare these intensities and positions to the expected values based on our forward models.

To perform this comparison, we determine the expected location of the diffraction peaks in camera coordinates, $\mathbf{r}_c = (x_c, y_c)$. We first map the pattern frequencies $\mathbf{f} = (f_x, f_y)$ to output angles characterized by the unit vectors $\hat{\mathbf{b}}(\mathbf{f})$, which according to eq. 6 are given by

$$b_{x,y}(\mathbf{f}) = b^o_{x,y} + \frac{\lambda}{d_{x,y}} f_{x,y}, \quad \textbf{(26)}$$

where the $\mathbf{f}$ are expressed in inverse number of mirrors and $\hat{\mathbf{b}}_o$ is the primary diffraction order of interest.

To map the output angles to pupil coordinates, we suppose the optical axis of the lens L6 is described by the unit vector $\hat{\mathbf{p}}$. To write the frequency in pupil coordinates, we must project the unit vectors $\hat{\mathbf{b}}(\mathbf{f})$ on the plane orthogonal to $\hat{\mathbf{p}}$. To achieve this, we construct an orthonormal coordinate system defined by $\hat{\mathbf{p}}$, $\hat{\mathbf{x}}_p = (p_z, 0, -p_x)/\sqrt{p_x^2 + p_z^2}$ and $\hat{\mathbf{y}}_p = \hat{\mathbf{p}} \times \hat{\mathbf{x}}_p$. Then the diffracted order positions in pupil coordinates are given by $\hat{\mathbf{x}}_p \cdot \hat{\mathbf{b}}(\mathbf{f})$ and $\hat{\mathbf{y}}_p \cdot \hat{\mathbf{b}}(\mathbf{f})$.

Finally, the camera coordinates are related to the pupil coordinates by an affine transformation with no shear

$$\begin{pmatrix} x_c(\mathbf{f}) \\ y_c(\mathbf{f}) \\ 1 \end{pmatrix} = M \begin{pmatrix} \cos\theta_c & -\epsilon \sin\theta_c & c_x \\ \sin\theta_c & \epsilon\cos\theta_c & c_y \\ 0 & 0 & 1 \end{pmatrix} \begin{pmatrix} \hat{\mathbf{x}}_p \cdot \hat{\mathbf{b}}(\mathbf{f}) \\ \hat{\mathbf{y}}_p \cdot \hat{\mathbf{b}}(\mathbf{f}) \\ 1 \end{pmatrix}, \quad \textbf{(27)}$$

where $\epsilon = \pm 1$ if the camera coordinates are not inverted (inverted) relative to the pupil coordinates and $(c_x, c_y)$ is the center of the pupil in camera coordinates. The magnification is the ratio of the lens focal length to the camera pixel size, $M = f_l/d_c$.

On the other hand, the intensities in the pupil can be obtained from eq. 5,

$$I(\mathbf{f}) \propto \text{sinc}^2\left[A_+\left(\gamma, \hat{\mathbf{a}} - \hat{\mathbf{b}}(\mathbf{f}), \hat{\mathbf{n}}\right)\right]\text{sinc}^2\left[A_-\left(\gamma, \hat{\mathbf{a}} - \hat{\mathbf{b}}(\mathbf{f}), \hat{\mathbf{n}}\right)\right] \quad (28)$$

$$A_+ = \begin{pmatrix} n_x^2(1-\cos\gamma)+\cos\gamma \\ n_x n_y(1-\cos\gamma)+n_z\sin\gamma \\ n_x n_z(1-\cos\gamma)-n_y\sin\gamma \end{pmatrix} \cdot \left(\hat{\mathbf{a}} - \hat{\mathbf{b}}\right) \quad (29)$$

$$A_- = \begin{pmatrix} n_x n_y(1-\cos\gamma)-n_z\sin\gamma \\ n_y^2(1-\cos\gamma)+\cos\gamma \\ n_y n_z(1-\cos\gamma)+n_x\sin\gamma \end{pmatrix} \cdot \left(\hat{\mathbf{a}} - \hat{\mathbf{b}}\right) \quad (30)$$

where we know emphasize the dependence of $A_\pm$ on the mirror rotation angle $\gamma$ and the rotation axis $\hat{\mathbf{n}}$, and provide expressions for the $A$'s for arbitrary $\hat{\mathbf{n}}$. In the main text, assume $\hat{\mathbf{n}} = (1,1,0)/\sqrt{2}$.
This provides us for a complete model of DMD diffraction. Now, we take wavelength $\lambda_i$ using diffraction orders $(n_i, -n_i)$, some of which rely on "on" mirror angles $\gamma_{\text{on}}$ and some which rely on the "off" mirror angles $\gamma_{\text{off}}$. Three sets of parameters must be determined: those related to the beam angles $\{b_x(\lambda_i), b_y(\lambda_i), \lambda_i\}$, the pupil mapping $\{M, \theta_c, p_x, p_y, c_x, c_y\}$, and the DMD parameters $\{\theta_n, d_x, d_y, w_x, w_y, \gamma_{\text{on}}, \gamma_{\text{off}}\}$. Here $\theta_n$ parameterizes the in-plane rotation of the DMD mirror rotation axis $\hat{\mathbf{n}}$, as we suppose $n_z = 0$. These are supplemented by the known values $\eta_i = \pm 1$ if the diffraction of interest comes from the "on" or "off" mirror state.
Many of these parameters are known to a reasonable degree of accuracy, but to find the best match of this model with the experimental data we perform a non-linear least squares fit using the cost function

$$\begin{aligned} C = & \sum_{f,i} \alpha \left( [x_c(\mathbf{f}) - x_c^e(\mathbf{f})]^2 + [y_c(\mathbf{f}) - y_c^e(\mathbf{f})]^2 \right) \\ & + \left( \frac{I^e(\mathbf{f})}{I^e(0)} / \left|\frac{E(\mathbf{f})}{E(0)}\right|^2 - \text{sinc}^2\left[A_+\left(\gamma_i, \hat{\mathbf{a}} - \hat{\mathbf{b}}_i(\mathbf{f}), \hat{\mathbf{n}}\right)\right]\text{sinc}^2\left[A_-\left(\gamma_i, \hat{\mathbf{a}} - \hat{\mathbf{b}}_i(\mathbf{f}), \hat{\mathbf{n}}\right)\right]\right)^2 \end{aligned} \quad (31)$$

where $\gamma_i = \gamma_{\text{on/off}}$ for $\eta_i = \pm 1$. Here the $E(\mathbf{f})$ are the predictions for the diffracted order strengths given by eq. 12. The parameter $\alpha$ weights the relative contributions of the position and intensity data and is set empirically to $1 \times 10^{-3}$, a value that achieves a reasonable balance between the position and amplitude cost functions. Changing $\alpha$ by an order of magnitude has a weak effect on the obtained parameter values.
Minimizing this cost function, we find excellent agreement between the theory and experimental results. The parameters obtained from this fit are shown in Fig. 1, and show excellent agreement with the expected values. A comparison between the experimental and fit results is provided in table 1. The high quality agreement between the experiment and theory results validates the models presented in the main text.

### 3: DMD diffraction efficiency

The DMD, like all binary diffractive optics, has relatively low diffraction efficiency, which requires use of high power lasers to illuminate a large FOV. Still, the blazed grating effect allows more efficient coupling into a single diffraction order compared with ferroelectric LCoS SLM's. We expect the peak diffraction efficiency using the all "on" pattern is about $0.91$, computed by summing the sinc envelope for all allowed diffraction orders. This efficiency is reduced to $\sim 0.69$ by reflections from the uncoated DMD window, the finite micromirror reflectivity, and the micromirror fill factor. When the DMD displays the SIM pattern, we expect a fraction of $0.076 \times 0.92 = 0.070$ of this power is diffracted into each of the $\mathbf{f} = \pm \mathbf{f}_o$ orders, where the first factor comes from the diffraction efficiency of the SIM pattern (i.e. eq. 12 in the main text), and the second from the Blaze envelope. This implies an overall efficiency of $\sim 0.097$. We confirm these estimates in the experiment, where we measure an efficiency of $0.60$ for the all "on" pattern using Gaussian beams with small waists compared to the DMD size. Using our typical experimental configuration, these efficiencies are further reduced by a factor of $\sim 2$ because the beam overfills the DMD aperture to create a more uniform illumination profile. In this configuration, we find the efficiency is $\sim 0.3$ for all "on" and $\sim 0.04$ and for the two SIM beams.
The efficiency of the light transfer is further limited by polarization effects and the finite efficiency of the polarization optics. Unpolarized light passing through the pizza polarizer reduces the efficiency by another factor of $0.5$, and the transmission efficiency of the polarizer reduces it by $0.5$–$0.9$, with

lower transmission in the blue and higher in the red. Thus, the net efficiency of the entire optical train is ∼0.01–0.02.

### 4: OTF measurement technique

The forward model for intensity at the imaging plane is the convolution of the electric field predicted by the DMD forward model band limited by the excitation imaging system optical transfer function. The projected intensity components are given by the autocorrelation of this signal, and the final imaged intensity is additionally blurred by the detection system OTF, $H(\mathbf{f})$

$$I(\mathbf{f}) = H(\mathbf{f}) \sum_{\mathbf{s}} \hat{\mathbf{e}}(\mathbf{s}) E(\mathbf{s}) \cdot \hat{\mathbf{e}}(\mathbf{s}-\mathbf{f}) E^*(\mathbf{s}-\mathbf{f}) \text{ circ}\left(\frac{|\mathbf{s}|}{f_{\text{max}}}\right) \text{ circ}\left(\frac{|\mathbf{s}-\mathbf{f}|}{f_{\text{max}}}\right) \tag{32}$$

$$= H(\mathbf{f}) \sum_{\mathbf{s}} \tilde{P}(\mathbf{s}) \tilde{P}^*(\mathbf{s}-\mathbf{f}) \hat{\mathbf{e}}(\mathbf{s}) \cdot \hat{\mathbf{e}}(\mathbf{s}-\mathbf{f}) \; H_{\text{DMD}}(\mathbf{s}) H^*_{\text{DMD}}(\mathbf{s}-\mathbf{f}) \text{ circ}\left(\frac{|\mathbf{s}|}{f_{\text{max}}}\right) \text{ circ}\left(\frac{|\mathbf{s}-\mathbf{f}|}{f_{\text{max}}}\right) \tag{33}$$

where $\mathbf{f}$ are restricted to the reciprocal lattice vectors of the projected pattern and the $P(\mathbf{f})$ can be obtained from eq. 12. When working with the $473\,\text{nm}$ excitation, the corrections from $H_{\text{DMD}}$ are $< 10\%$ out to the band limit. We neglect possible aberrations in the illumination pupil function, however these could be included in systems where they are relevant. The unit vectors $\hat{\mathbf{e}}(\mathbf{s})$ describe the polarization of the light after diffraction by the imaging system. This equation does not include the typical $\cos^{-1/2}\theta$ apodization factor that appears in the pupil of a system obeying the Abbe sine condition (4), due to the fact that this is a discrete sum and not an integral.
Neglecting $H_{\text{DMD}}$, we have

$$I(\mathbf{f}) \approx H(\mathbf{f}) \left[ \mathcal{O}(\mathbf{f}) \otimes \sum_{\mathbf{s}} \tilde{P}(\mathbf{s}) \tilde{P}^*(\mathbf{s}-\mathbf{f}) \hat{\mathbf{e}}(\mathbf{s}) \cdot \hat{\mathbf{e}}(\mathbf{s}-\mathbf{f}) \text{ circ}\left(\frac{|\mathbf{s}|}{f_{\text{max}}}\right) \text{ circ}\left(\frac{|\mathbf{s}-\mathbf{f}|}{f_{\text{max}}}\right) \right] \tag{34}$$

$$= H(\mathbf{f}) \left[ \mathcal{O}(\mathbf{f}) \otimes \sum_i m(\mathbf{f}_i) e^{i\phi(\mathbf{f}_i)} \delta(\mathbf{f}-\mathbf{f}_i) \right] \tag{35}$$

$$= \sum_i H(\mathbf{f}_i) m(\mathbf{f}_i) e^{i\phi(\mathbf{f}_i)} \mathcal{O}(\mathbf{f}-\mathbf{f}_i). \tag{36}$$

If we assume the object is perfectly uniform, this reduces to

$$I(\mathbf{f}_i) = \sum_i H(\mathbf{f}_i) m(\mathbf{f}_i) e^{i\phi(\mathbf{f}_i)} \delta(\mathbf{f}-\mathbf{f}_i) \tag{37}$$

$$H(\mathbf{f}_i) = \frac{m(\mathbf{f}_i) e^{i\phi(\mathbf{f}_i)}}{I(\mathbf{f}_i)}, \tag{38}$$

as given by eq. 1 in the main text.
To evaluate the quantities $m(\mathbf{f}_i)$, we must perform the sum above over the known Fourier components of the DMD pattern, and additionally evaluate the polarization vectors. We suppose the $\hat{\mathbf{z}}$ axis coincides with the optical axis, and define unit vectors and input polarization vectors by

$$\hat{\mathbf{e}}_p = (\cos\phi, \sin\phi, 0) \tag{39}$$

$$\hat{\mathbf{e}}_s = (-\sin\phi, \cos\phi, 0) \tag{40}$$

$$\hat{\mathbf{e}}_{\text{in}} = (\cos\phi_o, \sin\phi_o, 0). \tag{41}$$

After diffraction by the optical system, a ray initially incident parallel to the optical axis at azimuthal position $\phi$ converges towards the focal point and the polarization vector is modified (5)

$$\hat{\mathbf{e}}_{\text{out}}(\phi, \theta, \phi_o) = (\hat{\mathbf{e}}_p \cdot \hat{\mathbf{e}}_{\text{in}}) \hat{\mathbf{e}}_r + (\hat{\mathbf{e}}_s \cdot \hat{\mathbf{e}}_{\text{in}}) \hat{\mathbf{e}}_s \tag{42}$$

$$\hat{\mathbf{e}}_r = (\cos\phi\cos\theta, \sin\phi\cos\theta, \sin\theta). \tag{43}$$

In this measurement, we work with unpolarized light and hence we must average over the input polarization

$$\frac{1}{2\pi}\int_0^{2\pi} d\phi_o \hat{\mathbf{e}}(\phi_1,\theta_1,\phi_o)\cdot\hat{\mathbf{e}}(\phi_2,\theta_2,\phi_o) = \frac{1}{2}\cos^2(\phi_1-\phi_2)(1+\cos\theta_1\cos\theta_2) + \frac{1}{2}\cos(\phi_1-\phi_2)\sin\theta_1\sin\theta_2 + \frac{1}{2}\sin^2(\phi_1-\phi_2)(\cos\theta_1+\cos\theta_2). \quad (44)$$

Finally, we must convert between the angular representation used here and the spatial frequency representation discussed above. These are connected by

$$\mathbf{f} = \frac{n}{\lambda}(\cos\phi\sin\theta, \sin\phi\sin\theta) \quad (45)$$

In practice, the object fluorescence distribution is not perfectly uniform. This introduces correction terms caused by the same frequency mixing that SIM takes advantage of. The dominant term comes from the object spectrum itself, $H(\mathbf{f})\mathcal{O}(\mathbf{f})$. We can obtain this term directly from a widefield image and subtract it from the measured image. Performing higher order corrections is more challenging, as this requires knowledge of $m(\mathbf{f}_i)\mathcal{O}(\mathbf{f}-\mathbf{f}_i)H(\mathbf{f})e^{i\phi(\mathbf{f}_i)}$, which must be obtained through deconvolution and frequency shifting, similar to SIM reconstruction. To avoid these complications, we choose patterns where the higher order correction terms are unimportant. These patterns typically have large separation between different Fourier peaks, corresponding to large reciprocal lattice vectors and short periods. We find empirically that these corrections are not important for periods less than $\sim$20 mirrors.
To generate an OTF which is convenient to use with the SIM reconstruction algorithm, we empirically model the measured optical-transfer function as a Lorentzian times the ideal optical transfer function generated by the Airy point-spread function,

$$H(\mathbf{f}) = \frac{1}{1+\gamma^2|\mathbf{f}|^2}\times H_o(\mathbf{f}). \quad (46)$$

The Lorentzian attenuates the optical-transfer function at high frequencies. This functional form is convenient because it ensures that the OTF is unity at $\mathbf{f}=0$, zero at $|\mathbf{f}| = f_{\text{max}}$ and monotonically decreasing. The parameter $\gamma$, which controls the width of the Lorentzian, is determined by a non-linear least-squares fit.
This functional form assumes the OTF is azimuthally symmetric which is appropriate here because we find no evidence for a strong angular dependence in the experimental data. Nevertheless, a non-parametric OTF may be determined by smoothing and interpolating the data shown in Fig. 2F for systems which do not possess azimuthal symmetry.
Identifying the SIM peaks efficiently in the camera images requires careful calibration of the affine transformation between the DMD and the camera coordinates, which we obtain by projecting a test pattern of well-separated spots on the DMD, each generated by a single DMD micromirror. We perform Gaussian fits to these spots in the camera image. Additional image markers prevent ambiguities due to reflection and inversion. We then determine the affine transformation using Gaussian elimination.

### 5: SIM instrument design

A detailed description of the instrument optical path is given in the materials and methods section. The apparatus described there is shown in Fig. 3.

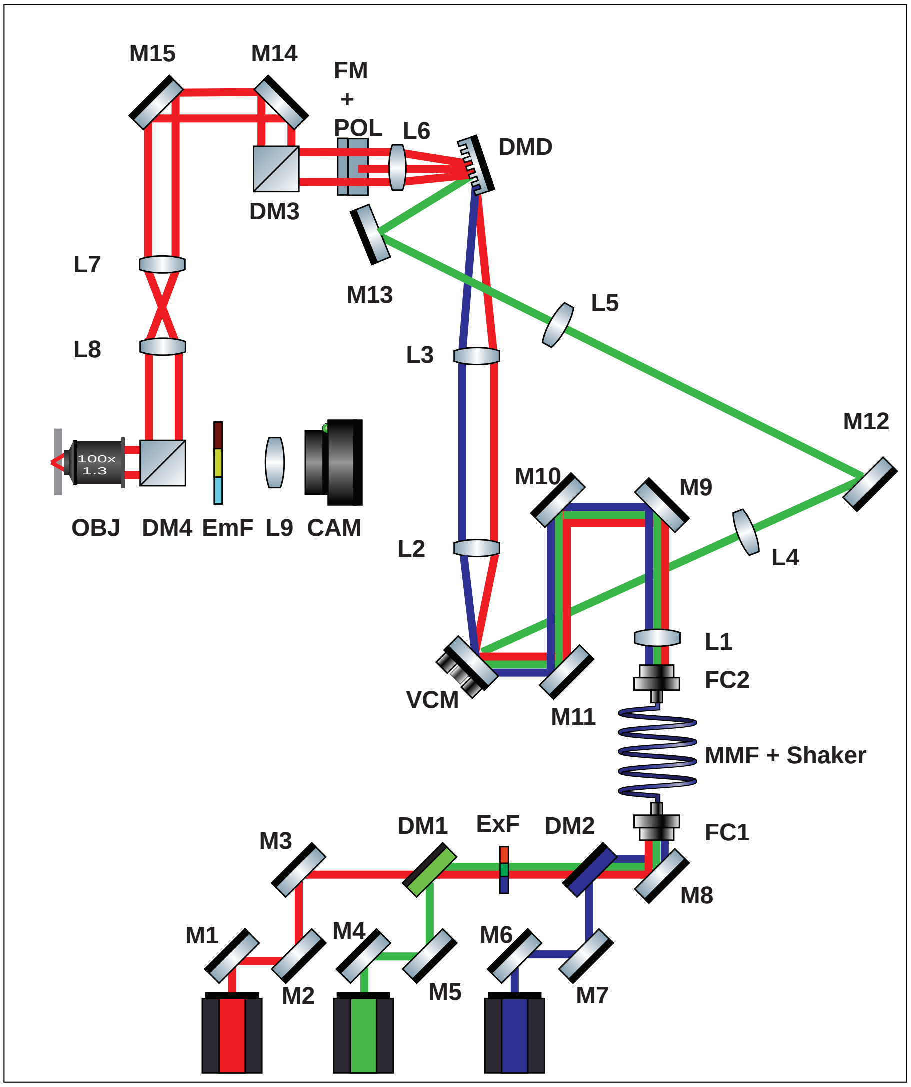

**Fig. 3. Schematic of microscope optical train.** Components of the three-color DMD microscope, corresponding to the parts described in table 2 of the main text.

### 6: Multicolor pattern generation algorithm

| wavelength | $\mathbf{r}_1$ | $\mathbf{r}_2$ | $P$ | $\theta$ |
|---|---|---|---|---|
| 473 nm | (-3, 11) | (3, 12) | 6.05 | 15.26° |
| | (-11, 3) | (12, 3) | 6.05 | 74.74° |
| | (-13, -12) | (12, 3) | 5.93 | 132.71° |
| 532 nm | (-3, 11) | (3, 15) | 6.84 | 15.26° |
| | (-11, 3) | (15, 3) | 6.84 | 74.74° |
| | (-13, -12) | (3, 12) | 6.78 | 132.71° |
| 635 nm | (-5, 18) | (-15, 24) | 8.03 | 15.52° |
| | (-18, 5) | (-24, 15) | 8.03 | 74.48° |
| | (-11, -10) | (15, 3) | 7.87 | 132.27° |

**Table 2.** Parameters of the DMD SIM patterns produced by the multicolor pattern generation algorithm.

We developed an algorithm to identify multicolor SIM patterns for an arbitrary number of wavelengths. The inputs to this algorithm are the approximate period of the desired SIM patterns, $P$, at one wavelength, the number of different angles, $n_a \geq 3$, the desired number of phases per pattern $n_p$ and a set of wavelengths $\lambda_i$. The criteria the patterns must satisfy are (1) the angles must be close to equally spaced by $\pi/n_a$, (2) the periods of patterns for different angles must match to good precision, (3) no parasitic Fourier modes from one pattern should overlap with the main frequency of another, and (4) patterns for different colors but the same angle must pass through the same location in the Fourier plane, which implies $P(\lambda_1)\lambda_1 = P(\lambda_2)\lambda_2$. Unlike previous implementations our pattern generation algorithm does not rely on exhaustive generation of lattice vectors (6).

Given a period $P$ and angle $\theta$, an algorithm to generate a single pattern must produce $\mathbf{r}_{1,2}$ such that $\mathbf{k}_2$ lies at angle $\theta$ and $1/|\mathbf{k}_2| = P$. Further, the imposed form of $\mathbf{r}_2 = (n,m)n_p$ constrains the reciprocal lattice vectors

$$\mathbf{k}_1 = \frac{1}{mr_{1x} - nr_{1y}}(m, -n) \tag{47}$$

$$\mathbf{k}_2 = \frac{1}{n_p(mr_{1x} - nr_{1y})}(r_{1y}, -r_{1x}). \tag{48}$$

Since the direction of $\mathbf{k}_1$ is already fixed, our strategy is to choose $1/|\mathbf{k}_2|$ close to the desired period $P$ with the constraint that the components of $\mathbf{r}_1$ must be integers. We must match both the angle and period of $\mathbf{k}_2$, i.e.

$$\theta = \arctan\left(-\frac{r_{1x}}{r_{1y}}\right) \tag{49}$$

$$P = 1/|\mathbf{k}_2| = n_p[n\cos(\theta) + m\sin(\theta)] \tag{50}$$

Our full algorithm generates the complete set of $n_a \times n_p$ patterns at each wavelength. We first describe the algorithm for generating a set of SIM patterns for one wavelength, and then discuss the generalization to multiple wavelengths. We begin by selecting a desired period, number of SIM angles $n_a$, and number of phase shifts $n_p$. Then we generate list of possible angles for integers $n, m < M$ using eq. 50. For each $n, m$ and fixed $P$, eq. 50 can be rewritten as a quadratic equation in $x = \cos\theta$. From this set of available angles we select sequences of angles $\theta_1 < \theta_2 < ... < \theta_{n_a}$, where the distance between adjacent angles is within a certain tolerance of the ideal spacing, $\pi/n_a$. We order our collection of sets of angles by the sizes of the $\mathbf{r}_2$ vectors,

$$C = \sum_{i=1}^{n_a} |\mathbf{r}_2(\theta_i)|. \tag{51}$$

This is a proxy for choosing a pattern set with well separated parasitic diffraction peaks, as large $\mathbf{r}_2$ vectors result in small reciprocal vectors.

For a given sequence of angles, we choose candidates for the vectors $\mathbf{r}_1$. For each angle, we generate $\mathbf{r}_1$ options by finding good rational approximations to $\tan\theta = -r_{1x}/r_{1y}$ for $r_{1x}, r_{2x} \in \{0, ..., N\}$. $N \approx 60$ is sufficient to match the desired $\theta$ to better than $1°$, which in turn ensures that the actual period $P$ matches the desired period to within a few percent.

We take all rational approximation of the angle within a certain tolerance, and select the set of $\mathbf{r}_1$ that maximize the distance between the SIM peaks and parasitic diffraction orders from other patterns. This is equivalent to maximizing the distance between $\mathbf{r}_2(\theta_i)$, the frequency of each SIM pattern, and all frequencies of each other pattern, which are given by $s\mathbf{k}_1(\theta_j)+t\mathbf{k}_2(\theta_j)$. This distance is given by

$$d_{\text{min}} = \min_{i\neq j}\left[\min_{s,t\in\mathbb{Z}}|\mathbf{k}_2(\theta_i)-s\mathbf{k}_1(\theta_j)-t\mathbf{k}_2(\theta_j)|\right]. \tag{52}$$

In practice, we only include reciprocal lattice vectors where the Fourier component of the pattern is nonzero in our minimization. For each angle set, we select the $r_1(\theta_i)$ by maximizing this distance. We select the final pattern by evaluating this for several angle sets.
This procedure is easily extended to multiple colors. When generating patterns for multiple wavelengths, we want to satisfy $P(\lambda_1)\lambda_1 = P(\lambda_2)\lambda_2$. In this case, we generate a set of angles for all wavelengths and specify an angular tolerance $\delta\theta$. For each angle, we check to ensure all wavelengths have an allowed angle within $\delta\theta$, and discard any angles that do not satisfy this constraint. Then, we proceed with the angle minimization routine described above, now taking the minimum pattern separation (scaled by the wavelength) over all colors. The results of this algorithm for a pattern period of ∼6 mirrors at $473\,\text{nm}$ are shown in Fig. 2.

### 7: SIM reconstruction

To mathematically express the image data obtained from a single raw SIM image with pattern having spatial frequency $\mathbf{f}_\theta$, angle $\theta$, and phase $\phi$, we use the following model

$$D_{\theta,\phi}(\mathbf{r}) = \left[P_{\theta,\phi}(\mathbf{r})O(\mathbf{r})\right]\otimes h(\mathbf{r})+N_{\theta,\phi} \tag{53}$$
$$P_{\theta,\phi} = A_{\theta\phi}\left[1+m_\theta\cos\left(2\pi\mathbf{r}\cdot\mathbf{f}_\theta+\phi\right)\right], \tag{54}$$

where the $A_{\theta\phi}$ are the amplitudes of the various patterns, the $m_\theta$ are the pattern modulation depths (which approach 1 in the ideal case), the $N_{\theta,\phi}$ are the measurement noise for each image, and $h$ is the (intensity) point-spread function of the detection imaging system.
In 2D SIM, 3 angles and 3 phases is sufficient to reconstruct the data naively. In this case, we can use the form of $P_{\theta,\phi}$ to simplify the forward model (eq. 53), if we neglect the noise terms. We find

$$\begin{pmatrix} D_{\theta,\phi_1}(\mathbf{f}) \\ D_{\theta,\phi_2}(\mathbf{f}) \\ D_{\theta,\phi_3}(\mathbf{f}) \end{pmatrix} = \begin{pmatrix} A_{\phi_1} & 0 & 0 \\ 0 & A_{\phi_2} & 0 \\ 0 & 0 & A_{\phi_3} \end{pmatrix} \begin{pmatrix} 1 & \frac{1}{2}e^{i(\phi_1-\phi_\theta^g)} & \frac{1}{2}e^{-i(\phi_1-\phi_\theta^g)} \\ 1 & \frac{1}{2}e^{i(\phi_2-\phi_\theta^g)} & \frac{1}{2}e^{-i(\phi_2-\phi_\theta^g)} \\ 1 & \frac{1}{2}e^{i(\phi_3-\phi_\theta^g)} & \frac{1}{2}e^{-i(\phi_3-\phi_\theta^g)} \end{pmatrix}$$
$$\times \begin{pmatrix} 1 & 0 & 0 \\ 0 & m_\theta e^{i\phi_\theta^g} & 0 \\ 0 & 0 & m_\theta s e^{-i\phi_\theta^g} \end{pmatrix} \begin{pmatrix} O(\mathbf{f})h(\mathbf{f}) \\ O(\mathbf{f}-\mathbf{f}_\theta)h(\mathbf{f}) \\ O(\mathbf{f}+\mathbf{f}_\theta)h(\mathbf{f}) \end{pmatrix}, \tag{55}$$

where we have factored out the modulation depth and a global phase offset $\phi_\theta^g$ to emphasize that we are free to regard these quantities as part of the transformation matrix or as part of object vector. From the latter point of view, only the phase differences are relevant for the transformation matrix.
To reconstruct the object $O(\mathbf{f})$ We take advantage of the known DMD patterns and the calibrated affine transformation between the DMD and camera planes to estimate the expected SIM patterns. We then estimate these using an optimization routine which determines the best pattern followed by independent routines to estimate the phase and modulation depth similar to (7).
Prior to parameter estimation, we preprocess the data by removing the camera background, estimated from the image histogram. We correct for intensity deviations between different phase images at each angle by matching the histograms using the scikit-image `match_histograms` function. This correction reduces reconstruction artifacts due to power fluctuations of our multimode laser sources.
Our conventional parameter estimation approach is similar to that described in (7). We use the known affine transformation from the DMD to camera space (8) to select a search region. We find an initial guess by taking the maximum of the cross correlation on a discrete set of points produced by a fast Fourier transform. We refine this selection, optimizing over continuous frequencies using the Fourier shift theorem. The phase and (optionally) amplitudes are estimated together using the correlation minimization method of Wicker (9). To obtain the modulation depth, we first fit the power spectrum

of the sample to an empirical power law form. Then we fit the region near the SIM peak to the same power law, but a different scaling constant. This constant is the modulation depth.
Once the SIM pattern parameters are determined, we invert eq. 55, obtaining

$$\begin{pmatrix} I_{\theta,0}(\mathbf{f}) \\ m_\theta e^{-i\phi_\theta^g} I_{\theta,1}(\mathbf{f}) \\ m_\theta e^{i\phi_\theta^g} I_{\theta,-1}(\mathbf{f}) \end{pmatrix} = M^{-1} \begin{pmatrix} D_{\theta,\phi_1}(\mathbf{f}) \\ D_{\theta,\phi_2}(\mathbf{f}) \\ D_{\theta,\phi_3}(\mathbf{f}) \end{pmatrix}, \quad \textbf{(56)}$$

where the $I_\epsilon(\mathbf{f})$ are essentially the $O(\mathbf{f}-\epsilon\mathbf{f}_\theta)h(\mathbf{f})$ up to the influence of noise. Before obtaining the $D(\mathbf{f})$ from $D(\mathbf{r})$ from a fast Fourier transform, we decompose $D(\mathbf{r})$ into a smooth and periodic part (10), and keep the periodic part. This is an alternative to multiplying $D(\mathbf{r})$ by an apodization function. We include a global phase factor in eq. 56, as in general there will be some error in our phase determination. We determine and correct this factor before the images are combined. The phase factor can be estimated based on the fact $|I_\theta(\mathbf{f})|$ should be real. Therefore, after shifting the components to the proper positions in Fourier space, we compute

$$\phi_\theta^g = -\text{angle}\left(\sum_{\mathbf{f}} \mathcal{T}_{\mathbf{f}_\theta}\left[m_\theta e^{-i\phi_\theta^g} I_{\theta,1}(\mathbf{f})\right] I_{\theta,0}^*(\mathbf{f})\right) \quad \textbf{(57)}$$

Finally, we reconstruct the object using a combination of Wiener deconvolution filters and weighted averaging

$$O(\mathbf{f}) = \sum_{\theta,\epsilon}\left[\frac{w_{\theta,\epsilon}(\mathbf{f}+\epsilon\mathbf{f}_\theta)}{\sum_{\theta,\epsilon} w_{\theta,\epsilon}(\mathbf{f}+\epsilon\mathbf{f}_\theta)+\eta}\right] e^{i\epsilon\phi_\theta^g}\mathcal{T}_{\epsilon\mathbf{f}_\theta}\left[\frac{h^*(\mathbf{f})}{|h(\mathbf{f})|^2\left(1+\frac{1}{w_{\theta,\epsilon}(\mathbf{f})}\right)} I_{\theta,\epsilon}(\mathbf{f})\right] A(\mathbf{f}), \quad \textbf{(58)}$$

where $\epsilon \in \{1, 0, -1\}$ and the weight factors are the signal-to-noise ratio, which is given by

$$w_{\theta,0}(\mathbf{f}) = |h(\mathbf{f})|^2 \frac{S_\theta(\mathbf{f})}{N_\theta} \quad \textbf{(59)}$$

$$w_{\theta,\epsilon=\pm 1}(\mathbf{f}) = m_\theta^2 |h(\mathbf{f})|^2 \frac{S_\theta(\mathbf{f}-\epsilon\mathbf{f}_\theta)}{N_\theta}. \quad \textbf{(60)}$$

Here $A$ is an apodization function, and $\eta > 0$ is a Wiener filter parameter that attenuates frequency points with low total weight. $\mathcal{T}_{\mathbf{f}_o} g(\mathbf{f}) = g(\mathbf{f}+\mathbf{f}_o)$ is the Fourier translation operator. The first bracketed term represents a weighted average, while the second is a Wiener filter deconvolution.
The signal-to-noise ratio is modeled using the phenomenological approach from (7). We assume the noise is white, and estimate the noise power by averaging the power spectral density beyond the optical transfer function cutoff frequency,

$$N_{\theta,\epsilon} = \int_{|\mathbf{f}|>f_{\text{max}}} \left|I_{\theta,\epsilon}(\mathbf{f})\right|^2. \quad \textbf{(61)}$$

We model the signal power spectral density as a power law, which is appropriate for many biological samples

$$S_\theta(\mathbf{f}) = A^2 |\mathbf{f}|^{-2\alpha}. \quad \textbf{(62)}$$

We restrict $0 \leq \alpha \leq 1.25$, which prevents observed bias towards large exponents in low SNR data.

### 8: Comparison of SIM reconstruction with FairSIM

To validate our SIM reconstruction algorithm we compare our results with FairSIM (11), a SIM reconstruction program written in Java which has been validated and is commonly used to reconstruct SIM data (11). We compare reconstructions of in-focus actin filaments imaged using the $473\,\text{nm}$ channel, which are presented in the main text Fig. 4C,D. Prior to performing the FairSIM reconstruction, we preprocess the raw SIM images using the histogram matching procedure described in section 7. We allow FairSIM to generate the OTF using $\text{na} = 1.3$, and attenuation factor $a = 0.3$. We perform the default FairSIM parameter finding routine, and the SIM frequencies it obtains agree with the expected values.

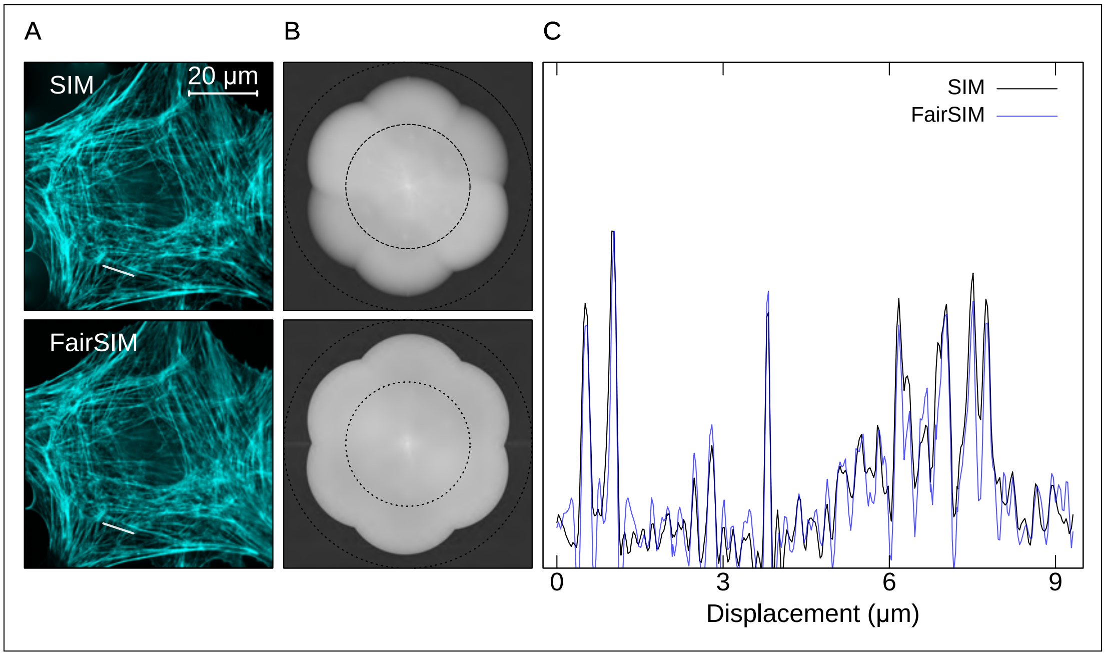

**Fig. 4. Comparison of reconstruction code with FairSIM. A**. SIM-SR reconstructions using our python reconstruction code (top) and FairSIM (bottom) for the in-focus slice of the BPAE cell images shown in Fig. 4C in the main text. The line cut shown in C. is illustrated by the semitransparent white line. **B**. Power spectral density of the images in A. **C**. Line cuts through our SIM-SR reconstruction (black) and the FairSIM reconstruction (blue). Each trace is normalized to its peak value. The two reconstruction techniques produce similar results.

We find our algorithm and FairSIM produce similar reconstructions, each showing similar increase in resolution and corresponding narrowing of filament width (Fig. 4). Because FairSIM uses a different normalization for their SR-SIM data, we first divide the line cut data by the peak value to allow easy comparison. All major features in the line cuts shown in Fig. 4C are present in both, although there are minor differences in smaller features. We also compare the resolution enhancements for the two approaches based on decorrelation analysis. We find that our reconstruction results in an enhancement of $1.54$, while theirs yields $1.75$ Both the differences in small features and in measured resolution enhancements are most likely due to the different approaches to Wiener filtering used in the two algorithms. These filters have different roll-off characteristics, resulting in different attenuation of high frequency SIM information. Furthermore, our algorithm applies a weighted averaging step which accounts for slightly different SNR's in the SIM images taken at different pattern angles. This effect is visible in the top panel of Fig. 4B, where the frequency information coming from the angle from the upper left to lower right corner is stronger than the other angles. In our reconstruction we use the experimental OTF, and in FairSIM we use their default OTF model. The close agreement in the reconstructed images despite these various implementation differences validates our approach.

### 9: SIM reconstruction of synthetic data

We reconstruct a variety of synthetic samples to further characterize our SIM algorithm and to assess the importance of using the experimentally obtained OTF versus the ideal model. We consider a set of variably spaced line pairs similar to the Argo-SIM slide and images of microtubules. Each of these samples show unique features in their Fourier space representations, making them ideal to test the SIM algorithm under a wide variety of conditions. For each sample, we perform reconstruction at a variety of signal-to-noise ratios. This is necessary to fully characterize the SIM reconstruction, as the maximum attainable resolution depends strongly on the SNR value.

In all cases, we first generate a ground truth image on an oversampled pixel grid. Next, we determine the fluorescence intensity by multiplying this with an illumination pattern and convolving with the point-spread function (PSF) generated from the experimental OTF. Then, we bin the oversampled grid to match the $\sim 60\,\mathrm{nm}$ pixel size in our imaging system. We normalize the image so that it has a certain peak photon number, and then we add Poisson noise to each pixel. Next we convert the pixels to camera counts by multiplying by a gain factor of $g = 2\,\mathrm{adu/e}$ and adding an offset of $100\,\mathrm{adu}$, and model camera readout noise by adding Gaussian noise with standard deviation of $\sigma = 4\,\mathrm{adu}$ to each pixel independently. This corresponds to noise of $2\,\mathrm{e}$, similar to the measured rms read noise of our sCMOS camera. We repeat this procedure for all phases and angles to produce synthetic raw SIM data. Finally, we apply the SIM reconstruction algorithm on this simulated image. For each image, we estimate the peak signal-to-noise as

$$\mathrm{SNR} = \frac{g p_{\mathrm{max}}}{\sqrt{g^2 p_{\mathrm{max}} + \sigma^2}}, \qquad \textbf{(63)}$$

where $p_{\mathrm{max}}$ is the peak average photon number incident on any pixel.

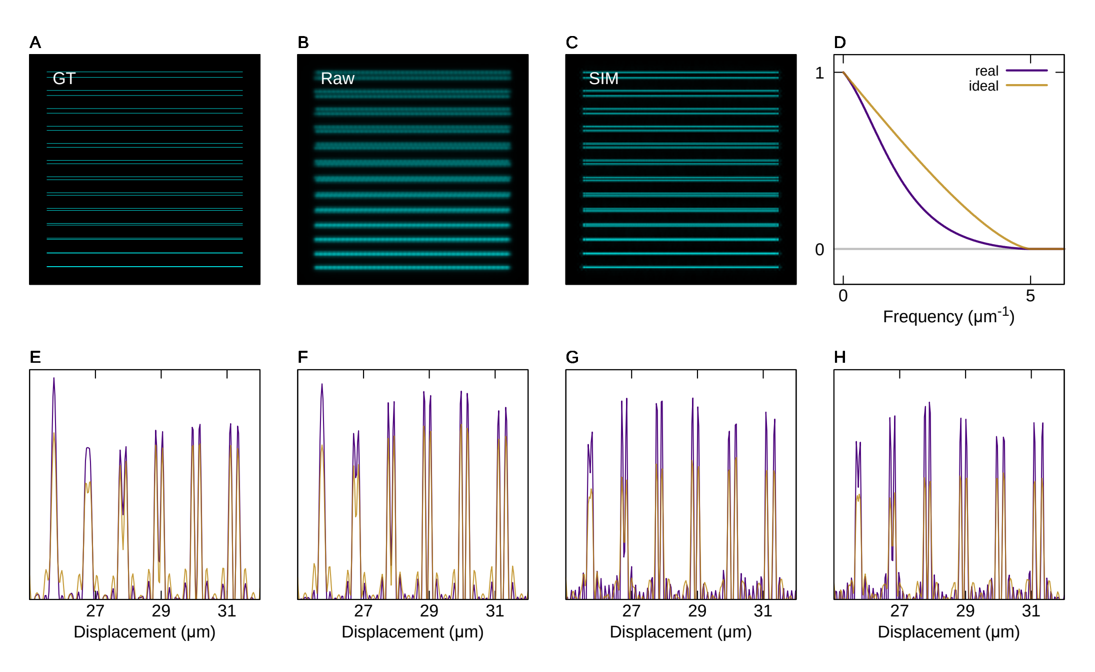

**Fig. 5. SIM Reconstruction of synthetic line pairs. A**. Ground truth image. Pixels are $15\,\mathrm{nm} \times 15\,\mathrm{nm}$, and the line pair separations range from $390\,\mathrm{nm}$ (top most) to $30\,\mathrm{nm}$ in steps of $30\,\mathrm{nm}$. **B**. A simulated SIM image with the same signal-to-noise ratio as H. **C**. SIM reconstruction using the experimental OTF for the same SNR as H. **D**. Optical transfer function used to generate the synthetic SIM images (purple) and ideal optical transfer function for $\mathrm{na} = 1.3$ and $\lambda = 520\,\mathrm{nm}$ (yellow). **E**. Line cut along the vertical direction showing lines with $90\,\mathrm{nm}$–$240\,\mathrm{nm}$. The peak photon number per pixel is $\sim$9000 photons, and peak signal-to-noise ratio is $\sim$30. **F**. As in D. but with SNR $\sim$90 **G**. SNR $\sim$300. **H**. SNR $\sim$1000.

**9.1. Synthetic line pairs.** We generate line pairs with separations ranging from $390\,\mathrm{nm}$ to $30\,\mathrm{nm}$ in steps of $30\,\mathrm{nm}$ on a $15\,\mathrm{nm} \times 15\,\mathrm{nm}$ pixel grid. We adjust the signal-to-noise ratio by changing the effective photon number. For each SNR, we reconstruct the synthetic raw SIM images (illustrated in Fig. 5B) using our reconstruction algorithm, once using the experimental OTF and once using the ideal OTF for the wavelength and numerical aperture (shown in Fig. 5D). This results in an SR image, as shown in Fig. 5C. We assess the quality of the reconstructions by comparing line cuts orthogonal to the line pairs in Fig. 5E–H.

The ultimate SIM resolution shows significant variation with signal-to-noise. Fig. 5E–H illustrates that, in this case, an SNR increase of a factor of $\sim$3 results in a resolution increase of about $30\,\mathrm{nm}$. At the highest SNR considered here, we saturate the expected SIM resolution. This demonstrates that too-small SNR can degrade SIM resolution in an experimental setting. Fig. 5 appears to show small splitting in the $90\,\mathrm{nm}$ line pair, which is smaller spacing than the ideal SIM resolution. This is an illustration of the difference between contrast and resolution.

We find that the experimental OTF modestly outperforms the ideal OTF, and the effect become more pronounced at larger SNR values. At lower SNR values, there are situations where the contrast between the smallest resolved line pair is slightly better for the ideal OTF (Fig. 5E,F). This is most likely an effect of the Wiener filtering, which will more strongly attenuate high frequency information for the experimental OTF, which has less weight at these frequencies. On the other hand, the experimental OTF will correctly weight these frequencies during Wiener deconvolution when the SNR is large enough for the Wiener filter attenuation to be negligible (Fig. 5G,H).

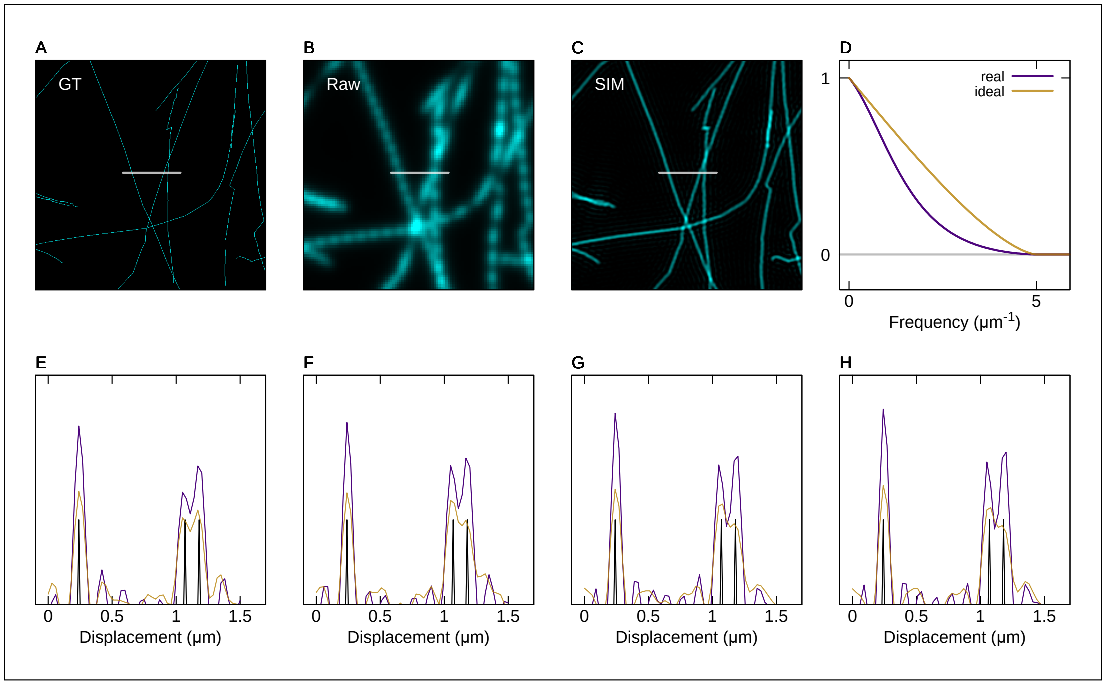

**Fig. 6. SIM Reconstruction of synthetic microtubules. A**. Ground truth image. Pixels are $10\,\mathrm{nm} \times 10\,\mathrm{nm}$, and the ground truth structures have width of $10\,\mathrm{nm}$. The white line illustrates the location of the line cuts shown in E.-H. **B**. A simulated SIM image with the same signal-to-noise ratio as H. **C**. SIM reconstruction using the experimental OTF for the same SNR as H. **D**. Optical transfer function used to generate the synthetic SIM images (purple) and ideal optical transfer function for $\mathrm{na} = 1.3$ and $\lambda = 520\,\mathrm{nm}$ (yellow). **E**. Line cut showing one isolated microtubule (left peak) and two microtubules near a crossing (right peak). The ground truth is shown in black, the SIM reconstruction using the experimental OTF is shown in purple, and the reconstruction using the ideal OTF is shown in yellow. The peak photon number per pixel is $\sim 4000$ photons, and peak signal-to-noise ratio is $\sim 60$. **F**. As in D. but with SNR $\sim 200$ **G**. SNR $\sim 650$. **H**. SNR $\sim 2000$.

**9.2. Synthetic microtubules.** We generate a ground truth image of 79 one-pixel thick lines representing microtubules on a $10\,\mathrm{nm} \times 10\,\mathrm{nm}$ pixel grid with full size $20.46\,\mathrm{\mu m} \times 20.46\,\mathrm{\mu m}$. The lines are generated from a filament network file provided by Super Resolution Simulation (SuReSim) (12). For simplicity, we ignore the $z$-coordinate and assume the microtubules are all in the same plane. We illustrate a $6\,\mathrm{\mu m} \times 6\,\mathrm{\mu m}$ portion of the ground truth, simulated SIM image for one angle and phase, and SIM reconstruction in Fig. 6A, B, and C respectively. We assess the SIM algorithm performance by looking at a line cut through the reconstructed data near a crossing between two microtubules, where the microtubules are separated by $110\,\mathrm{nm}$ (Fig. 6).

We find that the SIM reconstruction using the experimental OTF can distinguish the closely spaced microtubules for sufficiently large SNR, but the reconstruction using the ideal OTF cannot (Fig. 6E-H). The similarity between Fig. 6G and H indicates that the reconstruction quality of these two images is no longer limited by SNR. The lower resolution of the ideal OTF reconstruction is expected because the Wiener deconvolution does not correctly reweight the high frequency information in Fourier space. This demonstrates that using an experimentally determined OTF can provide quantitative and qualitative improvement over the ideal OTF.

### 10: Comparison of coherent and incoherent-projection SIM

SIM using a DMD and *incoherent* projection has previously been realized, but while this technique has the same theoretical resolution as coherent SIM, its resolution in the presence of noise degrades much more rapidly than coherent SIM. The achievable signal in incoherent projection SIM is smaller than coherent SIM by a factor of the optical transfer function at the SIM frequency, $H(\mathbf{f}_o)$. Because the experimental resolution limit of SIM is set by the signal-to-noise in the image, coherent SIM has a tremendous advantage over incoherent projection techniques. Achieving comparable results with incoherent projection SIM requires at least one order of magnitude more optical power for realistic SIM frequencies, making it much less efficient with its photon budget and leading to higher photobleaching and phototoxicity.

Coherent SIM relies on interfering two beams which pass near the edge of the objective pupil. Assuming an ideal optical transfer function, these beams are not attenuated by the OTF, leading to a

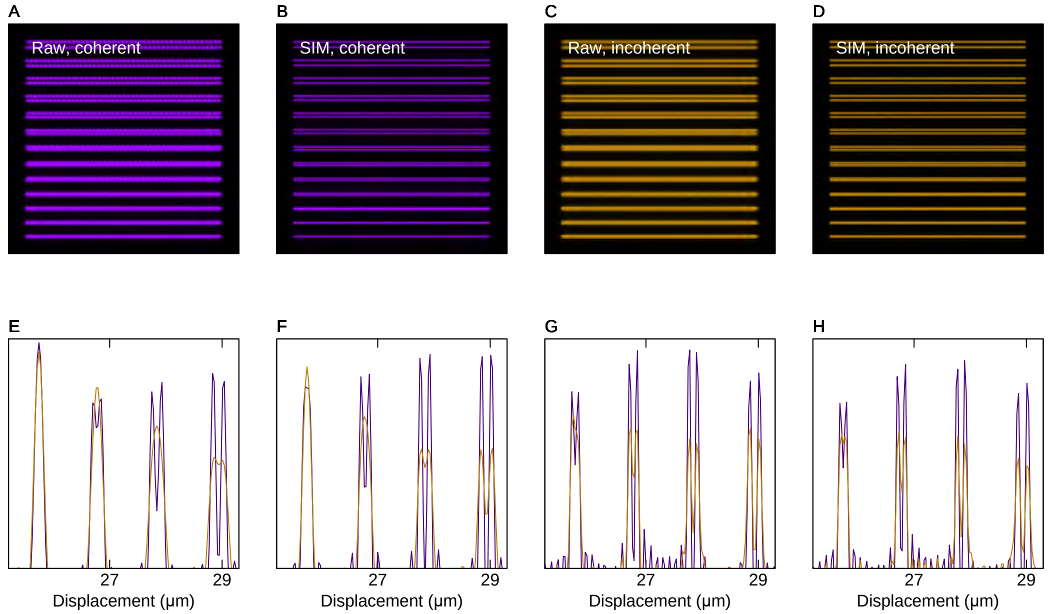

**Fig. 7. Comparison of coherent and incoherent SIM versus SNR. A**. Simulated raw image for one angle and phase combination using coherent SIM. The Moiré pattern features are clearly visible. The SNR in this image is the same as in G. **B**. Reconstructed SIM image corresponding to the raw data in A. **C**. Simulated raw image for one angle and phase combination using incoherent SIM. The Moiré pattern is too weak to be seen by eye. The SNR in this images is the same as in G. **D**. Reconstructed SIM image corresponding to the raw data in C. **E**. Line cut illustrating coherent (magenta) and incoherent (yellow) SIM for fixed net intensity at the sample and SNR $\sim 30$, corresponding to peak photon number of $\sim 1000$ photons per pixel. Lines with spacings ranging from $90\,\mathrm{nm}$ to $180\,\mathrm{nm}$ are shown. The At this SNR, the largest three line pairs are resolved by coherent SIM, but not by incoherent SIM. **F**. SNR $\sim 100$. Coherent SIM clearly resolves the first three peaks. Incoherent SIM resolves the first peak, but with worse contrast that coherent SIM. **G**. SNR $\sim 300$. Coherent SIM resolves all four peaks, while incoherent SIM resolves only the first three but with significantly less contrast. **H**. SNR $\sim 1000$. Despite increased signal to noise, the reconstructed SIM images are similar to G. Increasing signal to noise further has only a marginal effect.

sinusoidal intensity pattern with full contrast ($m = 1$),

$$I_c^{\text{sample}} = I_o \left(1 + \cos\left(2\pi \mathbf{f}_o \cdot \mathbf{r} + \phi\right)\right). \tag{64}$$

Incoherent SIM, on the other hand, relies on forming an image of a sinusoidal *intensity pattern*, and the strength of the modulation is degraded by the optical transfer function

$$I_i^{\text{sample}} = I_o \left(1 + H(\mathbf{f}_o) \cos\left(2\pi \mathbf{f}_o \cdot \mathbf{r} + \phi\right)\right), \tag{65}$$

implying the modulation depth can be at most $m = H(\mathbf{f}_o)$.

The practical resolution of a SIM measurement is set by the SNR, and the strength of the signal containing superresolution information is determined by $I_o m$. For fixed intensity $I_o$, achieving the same $I_o m$ value requires a factor of $1/H(\mathbf{f}_o)$ power in the incoherent case. This lower efficiency also leads to enhanced photobleaching and phototoxicity.

Assuming an ideal optical transfer function, setting the SIM pattern at $80\,\%$ of the band pass limits the incoherent projection $m < 0.1$. Increasing the SIM frequency further to achieve the full SIM theoretical resolution limit causes the modulation to rapidly deteriorate. Setting the SIM pattern at $95\,\%$ of the band pass limits $m < 0.01$, implying that coherent SIM can achieve SNR's a factor of $100\times$ larger than with an equivalent incoherent system. This value is similar to that chosen in (13), an experiment that would not have been possible using an incoherent projection SIM.

These numbers provide a lower bound on the SNR advantage coherent SIM has over incoherent projection because experimental optical transfer functions typically decay faster than the ideal OTF. The experimental OTF we obtained in section 2.1 implies $m < 0.02$ and $0.002$ at $80\,\%$ and $95\,\%$ of the theoretical band pass, respectively.

To illustrate the effect this difference has on the image quality, we simulated SIM raw images and reconstructions of the same line pairs discussed in section 9. We summarize the results in Fig. 7. We find that coherent SIM resolves smaller line pairs and thus achieves significantly better resolution than incoherent SIM at all SNR values we consider (Fig. 7E-H). As expected, the weaker modulation in the incoherent case leads to a visibly weaker Moiré pattern in the raw SIM images (Fig. 7A,C).

### 11: Additional SIM experimental tests

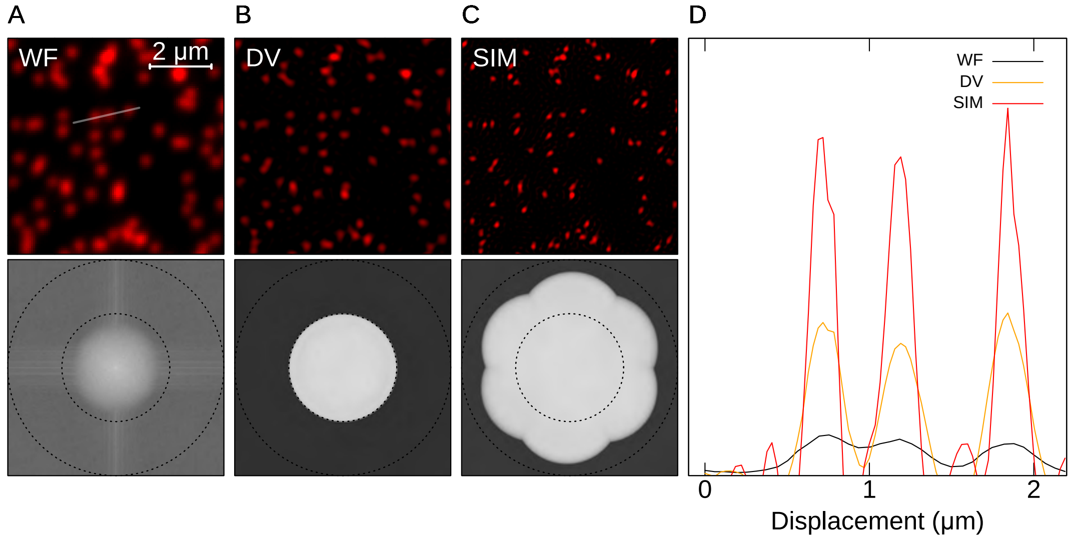

**Fig. 8. SIM performance on diffraction limited spots. A**. Widefield image for $635\,\mathrm{nm}$ (top) and power spectrum (bottom). Images are shown in ADU. Circles illustrate the maximum frequency where the ideal optical transfer function has support for $\lambda_{\text{emission}} = 680\,\mathrm{nm}$ and $\mathrm{na} = 1.3$. **B**. Widefield image after Wiener filter deconvolution. **C**. SR-SIM images. **D**. One-dimensional cuts plotted along the lines illustrated in A. We show the widefield image (black line), Wiener deconvolved image (orange), and SR-SIM image (red).

**11.1. Gattaquant DNA origami nanorulers.** We further assessed the SIM performance on a collection of diffraction limited objects, using Gattaquant DNA origami nanorulers. The nanorulers contain two dye molecules separated by a well defined distance of $120\,\mathrm{nm}$. They are designed for use in characterizing STED systems, and we did not expect to resolve this small spacing using the $680\,\mathrm{nm}$ fluorescence where the maximum theoretical resolution is $\sim 130\,\mathrm{nm}$. However, the very different characteristics of this sample compared with the fixed cells allowed an independent test of our SIM approach. In these samples the SIM peaks appear weaker due to the smaller DC spatial frequency component as compared with cell samples. However, high frequency content is still efficiently mixed below the band pass, resulting in a large SNR and high-quality reconstruction. As illustrated in Fig. 8, the SIM images show significant narrowing of the individual origami as compared with both the widefield images and deconvolved images. Decorrelation analysis estimates SIM provides a factor of 1.64 resolution enhancement, close to the theoretical maximum based on our pattern spacing. Performing Gaussian fits to $\sim$2500 DNA origami and comparing the resulting standard deviations for the deconvolved and SIM images gives a similar resolution enhancement estimate of 1.71.

**11.2. Three-wavelength imaging of fixed cells.** We realized three-wavelength SIM using the $473\,\mathrm{nm}$, $532\,\mathrm{nm}$, and $635\,\mathrm{nm}$ channels to image $\beta$-actin, HIF, and FGF18 labeled with Alexa Fluor 488, Alexa Fluor 555, and Alexa Fluor 647 in fixed primary rat endothelial cells. We have previously studied angiogenic signaling in primary rat, sheep, and human endothelial cells using high-throughput multi-wavelength widefield microscopy (14–16).

The SIM results (Fig. 9) show resolution enhancement in all three channels, with the enhancement in the details of the HIF distribution from the $532\,\mathrm{nm}$ channel appearing most visible in Fig. 9C. Decorrelation analysis estimates the SIM images enhance the resolution over the deconvolved images by by factors of $\sim$1.6, $\sim$1.5, and $\sim$1.65 for the $473\,\mathrm{nm}$, $532\,\mathrm{nm}$, and $635\,\mathrm{nm}$ channels respectively.

Primary rat endothelial cells were isolated as previously described (16, 17). After isolation, cells were plated on glass chamber slides (Thermo Fisher, 177402), fixed using $4\,\%$ paraformaldehyde (Electron Microscopy Sciences, 15710-S), and labeled using primary and secondary antibodies for HIF (Primary, 1:250 dilution: Thermo Fisher PA1-184. Secondary, 1:500 dilution, Alexa Fluor 555: Thermo Fisher A32794), $\beta$-actin (Primary, 1:1000 dilution: Thermo Fisher AM4302. Secondary, 1:500 dilution, Alexa Fluor 488: Thermo Fisher A-11001), and FGF18 (Primary, 1:250 dilution: Santa Cruz SC-16830. Secondary, 1:500 dilution, Alexa Fluor 647: Thermo Fisher A32849)). After labeling, slides were mounted with Slowfade Glass with DAPI (Thermo Fisher S36920) using $25\,\mathrm{mm} \times 40\,\mathrm{mm}$ 1.5 coverslips (Thermo Fisher 24x40-1.5).

1. James E. Harvey and Richard N. Pfisterer. Understanding diffraction grating behavior: including conical diffraction and Rayleigh anomalies from transmission gratings. *Optical Engineering*, 58(08):1, aug 2019. doi: 10.1117/1.oe.58.8.087105.

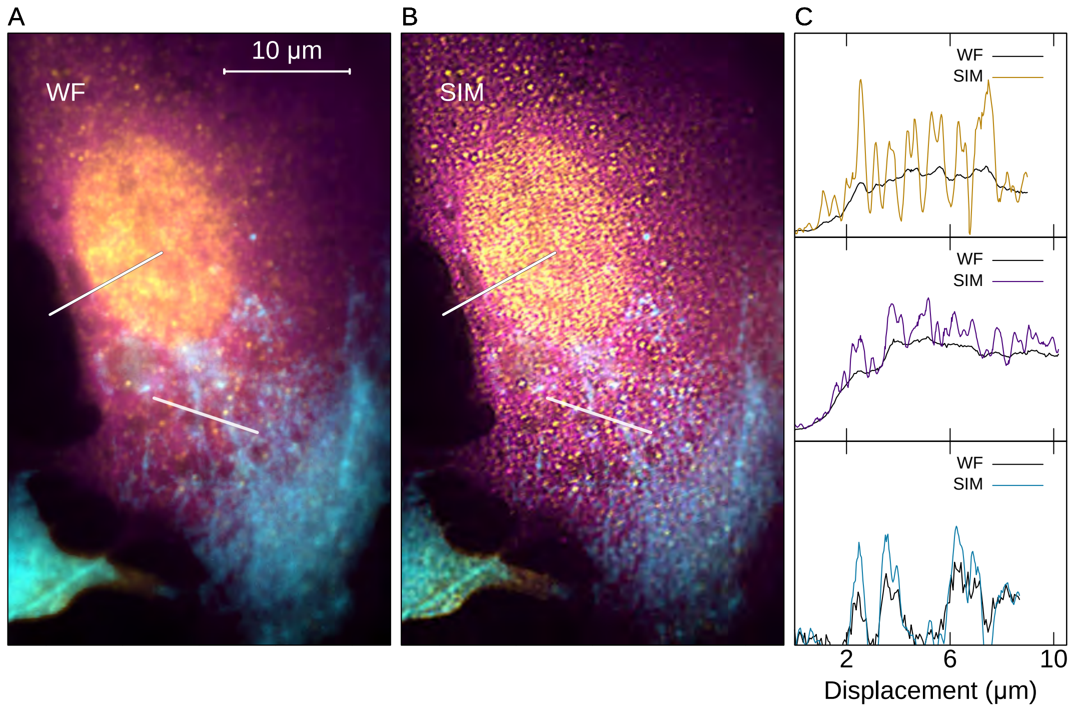

**Fig. 9. Three-color imaging of fixed primary rat endothelial cells. A**. Composite widefield image with the $473\,\mathrm{nm}$ channel (cyan), $532\,\mathrm{nm}$ channel (yellow), and $635\,\mathrm{nm}$ channel (magenta). Images are shown in ADU. White lines illustrate the position of the line cut shown in C. **B**. SR-SIM images. **C**. One-dimensional cuts plotted along the lines illustrated in A. for $532\,\mathrm{nm}$ (top), $635\,\mathrm{nm}$ (middle), and $473\,\mathrm{nm}$ excitations. $473\,\mathrm{nm}$ and $532\,\mathrm{nm}$ channel line cuts correspond to the upper line cut in A., and $635\,\mathrm{nm}$ to the lower cut.

2. Alice Sandmeyer, Mario Lachetta, Hauke Sandmeyer, Wolfgang Hübner, Thomas Huser, and Marcel Müller. DMD-based super-resolution structured illumination microscopy visualizes live cell dynamics at high speed and low cost. 2019.
3. Mario Lachetta, Hauke Sandmeyer, Alice Sandmeyer, Jan Schulte am Esch, Thomas Huser, and Marcel Müller. Simulating digital micromirror devices for patterning coherent excitation light in structured illumination microscopy. oct 2020.
4. Marcel Leutenegger, Ramachandra Rao, Rainer A. Leitgeb, and Theo Lasser. Fast focus field calculations. *Optics Express*, 14(23):11277, 2006. doi: 10.1364/oe.14.011277.
5. B. Richards and E. Wolf. Electromagnetic diffraction in optical systems, II. structure of the image field in an aplanatic system. *Proceedings of the Royal Society of London. Series A. Mathematical and Physical Sciences*, 253(1274):358–379, dec 1959. doi: 10.1098/rspa.1959.0200.
6. Hui-Wen Lu-Walther, Martin Kielhorn, Ronny Förster, Aurélie Jost, Kai Wicker, and Rainer Heintzmann. fastSIM: a practical implementation of fast structured illumination microscopy. *Methods and Applications in Fluorescence*, 3(1):014001, jan 2015. doi: 10.1088/2050-6120/3/1/014001.
7. Amit Lal, Chunyan Shan, and Peng Xi. Structured illumination microscopy image reconstruction algorithm. *IEEE Journal of Selected Topics in Quantum Electronics*, 22(4):50–63, jul 2016. doi: 10.1109/jstqe.2016.2521542.
8. Pavel Křížek, Ivan Raška, and Guy M. Hagen. Flexible structured illumination microscope with a programmable illumination array. *Optics Express*, 20(22): 24585, oct 2012. doi: 10.1364/oe.20.024585.
9. Kai Wicker, Ondrej Mandula, Gerrit Best, Reto Fiolka, and Rainer Heintzmann. Phase optimisation for structured illumination microscopy. *Optics Express*, 21(2):2032, jan 2013. doi: 10.1364/oe.21.002032.
10. Lionel Moisan. Periodic plus smooth image decomposition. *Journal of Mathematical Imaging and Vision*, 39(2):161–179, oct 2010. doi: 10.1007/s10851-010-0227-1.
11. Marcel Müller, Viola Mönkemöller, Simon Hennig, Wolfgang Hübner, and Thomas Huser. Open-source image reconstruction of super-resolution structured illumination microscopy data in ImageJ. *Nature Communications*, 7(1), mar 2016. doi: 10.1038/ncomms10980.
12. Varun Venkataramani, Frank Herrmannsdörfer, Mike Heilemann, and Thomas Kuner. SuReSim: simulating localization microscopy experiments from ground truth models. *Nature Methods*, 13(4):319–321, feb 2016. doi: 10.1038/nmeth.3775.
13. Loïc Reymond, Johannes Ziegler, Christian Knapp, Fung-Chen Wang, Thomas Huser, Verena Ruprecht, and Stefan Wieser. SIMPLE: Structured illumination based point localization estimator with enhanced precision. *Optics Express*, 27(17):24578, aug 2019. doi: 10.1364/oe.27.024578.
14. Megan A Ahern, Claudine P Black, Gregory J Seedorf, Christopher D Baker, and Douglas P Shepherd. Hyperoxia impairs pro-angiogenic RNA production in preterm endothelial colony-forming cells. *AIMS Biophysics*, 4(2):284–297, 2017. ISSN 2377-9098. doi: 10.3934/biophy.2017.2.284.
15. Christina Kim, Gregory J Seedorf, Steven H Abman, and Douglas P Shepherd. Heterogeneous response of endothelial cells to insulin-like growth factor 1 treatment is explained by spatially clustered sub-populations. *Biology Open*, 8(11), November 2019. ISSN 2046-6390. doi: 10.1242/bio.045906.
16. Gregory Seedorf, Christina Kim, Bradley Wallace, Erica W Mandell, Taylor Nowlin, Douglas Shepherd, and Steven H Abman. rhIGF-1/BP3 preserves lung growth and prevents pulmonary hypertension in experimental bronchopulmonary dysplasia. *American Journal of Respiratory and Critical Care Medicine*, 201 (9):1120–1134, May 2020. ISSN 1073-449X, 1535-4970. doi: 10.1164/rccm.201910-1975OC.
17. Cristiana Iosef, Tero-Pekka Alastalo, Yanli Hou, Chihhsin Chen, Eloa S Adams, Shu-Chen Lyu, David N Cornfield, and Cristina M Alvira. Inhibiting nf-$\kappa$b in the developing lung disrupts angiogenesis and alveolarization. *American journal of physiology. Lung cellular and molecular physiology*, 302(10):L1023–36, May 2012. ISSN 1040-0605, 1522-1504. doi: 10.1152/ajplung.00230.2011.
